# Optimization algorithm for omic data subspace clustering

**DOI:** 10.1101/2021.11.12.468415

**Authors:** Madalina Ciortan, Matthieu Defrance

**Author notes:** Permission to make digital or hard copies of all or part of this work for personal or classroom use is granted without fee provided that copies are not made or distributed for profit or commercial advantage and that copies bear this notice and the full citation on the first page. Copyrights for components of this work owned by others than ACM must be honored. Abstracting with credit is permitted. To copy otherwise, or republish, to post on servers or to redistribute to lists, requires prior specific permission and/or a fee. Request permissions from.

## Abstract

Subspace clustering identifies multiple feature subspaces embedded in a dataset together with the underlying sample clusters. When applied to omic data, subspace clustering is a challenging task, as additional problems have to be addressed: the curse of dimensionality, the imperfect data quality and cluster separation, the presence of multiple subspaces representative of divergent views of the dataset, and the lack of consensus on the best clustering method.

First, we propose a computational method (*discover*) to perform subspace clustering on tabular high dimensional data by maximizing the internal clustering score (i.e. cluster compactness) of feature subspaces. Our algorithm can be used in both unsupervised and semi-supervised settings. Secondly, by applying our method to a large set of omic datasets (i.e. microarray, bulk RNA-seq, scRNA-seq), we show that the subspace corresponding to the provided ground truth annotations is rarely the most compact one, as assumed by the methods maximizing the internal quality of clusters. Our results highlight the difficulty of fully validating subspace clusters (justified by the lack of feature annotations). Tested on identifying the ground-truth subspace, our method compared favorably with competing techniques on all datasets. Finally, we propose a suite of techniques to interpret the clustering results biologically in the absence of annotations. We demonstrate that subspace clustering can provide biologically meaningful sample-wise and feature-wise information, typically missed by traditional methods.

CCS Concepts: • **Computing methodologies** → **Genetic algorithms**; **Mixture models**; **Cluster analysis**.

**ACM Reference Format:** Madalina Ciortan and Matthieu Defrance. 2021. Optimization algorithm for omic data subspace clustering. 1, 1 (September 2021), 40 pages. https://doi.org/10.1145/nnnnnnn.nnnnnnn

## 1 INTRODUCTION

Subspace clustering studies the techniques to partition datasets feature-wise and sample-wise. Applied to omic datasets, subspace clustering makes possible the direct identification of both the set of transcripts/genes (*S*_*i*_ in Figure 1a) and the underlying groups of patients/cells (*S*_*ij*_ Figure 1a). In contrast with subspace clustering, traditional clustering methods rely on the entire dataset to produce only one clustering solution. In the omic context, the salient features (i.e. genes) corresponding to sample clusters are as important as the identified clusters for the downstream analysis.

**Fig. 1.**
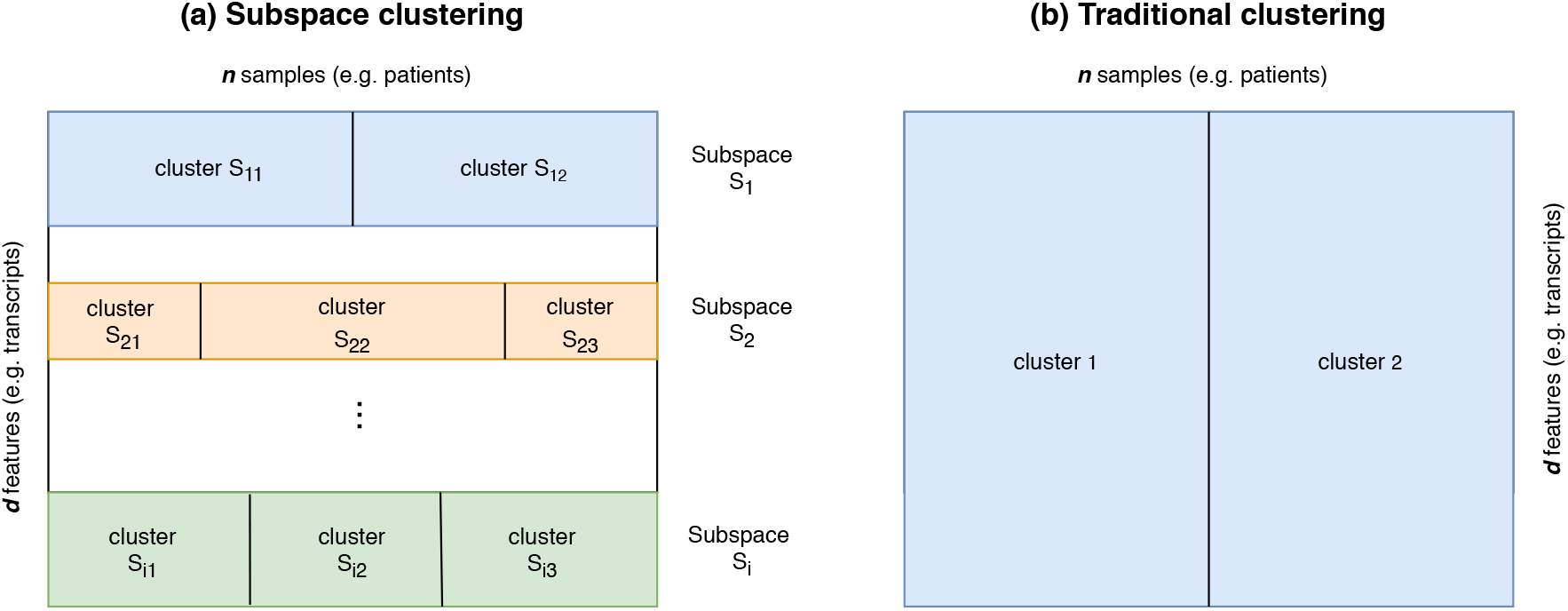
Subspace clustering compared to traditional clustering. Subspace clustering (a) identifies multiple subspaces (*S*_1_, *S*_2_ and *S*_*i*_) with distinct sample partitions (e.g. cluster *S*_11_ and *S*_12_ for subspace *S*_1_). In traditional clustering (b), all features are used to identify only one global sample partition (cluster 1 and cluster 2).

Parsons et al. [46] classified subspace clustering methods as bottom-up or top-down. Bottom-up approaches build up clusters iteratively, starting from dense units found in low dimensional subspaces. First publications leveraged either static grids (Clique [2], Encluse [9]) or adaptive grids (CLtree [37]). Top-down methods start with an initial approximation of the clusters in the full features space, followed by multiple iterations to refine the clusters. Other categories of subspace clustering algorithms leverage factorization-based algebraic [15, 54], statistical approaches [4] as well as spectral [18] or evolutionary [56] techniques.

Biclustering methods such as Qubic [16] build a graph where genes are connected with edges weighted by their co-expression rate. Qubic identifies biclusters corresponding to heavy sub-graphs, but does not impose conditions on the number or compactness of sample clusters. In our work, subspace clustering assumes that relevant subspaces contain at least 2 compact clusters and subspace clusters all samples.

The analysis of omic datasets brings computational challenges consisting of the high number of dimensions, the complex feature dependencies (e.g. gene co-expression) and the presence of noise. This motivated the proposal of numerous dedicated methods [30, 38, 55]. Most techniques propose performing successive steps, starting with filtering out unreliable observations from the dataset, normalizing the remaining values, applying feature selection methods and/or a dimensionality reduction technique to mitigate ‘the curse of dimensionality’. Some methods [33, 58] create a distance matrix from the low-dimensional representation of the dataset (usual choices of distance being euclidean, mutual information, Pearson and Spearman correlations [22, 29]). The distance matrix is then either used to construct a graph or integrated directly in a clustering algorithm (commonly KMeans [12, 32, 53], hierarchical clustering [59] and density-based clustering [47]). The high dimensionality can also be addressed with methods such as filter feature selection which evaluate various statistical properties of features and is usually computationally efficient [19].

Single-cell RNA-seq (scRNA-seq) data provides transcriptomic profiling for individual cells and introduces the dropout (i.e. false zero counts observations) as an additional computational challenge. Several methods were proposed to analyze scRNA-seq data. ScRNA [41] employs non-negative matrix factorization to incorporate information from a larger annotated dataset and applies transfer learning to perform the clustering. SOUP [63] handles both pure and transitional cells; it uses the expression similarity matrix to produce soft cluster memberships. Seurat [49] leverages graph-processing techniques like community detection (i.e. the Louvain algorithm) to produce a shared nearest neighbor graph and then to predict cluster assignments. RaceID [27] identifies rare cell types and improves clustering performance using K-medoids (instead of the traditional KMeans).

Despite the abundance of existing clustering methods, there is no consensus on the best approach. It was showed [21] that while the clustering solutions of various methods appear robust when analyzing the same datasets, the solutions have little in common with each other or with the supervised labeling. Moreover, despite the numerous proposed metrics for assessing the internal quality of clustering [48], the field has not not converge to a unified approach.

## 2 METHOD

We propose *discover*, an optimization algorithm performing bottom-up subspace clustering on tabular data in order to identify the subspaces with the most compact sample clusters. Given a dataset *D* = {*x*_*i j*_} ∈ ℝ^*d*\**n*^ with *d* features (i.e. transcripts) and *n* samples (i.e. patients), *discover* identifies a set of subspaces (i.e. the group of features *S*_*i*_ = {*x*_1_, · · · *x*_*m*_}, *m* < *d*) and the corresponding sample clusters, such that the partitioning (i.e. the sample clusters) of the subspace has maximal internal clustering score (compactness). Instead of defining a lower bound for the desired internal clustering score, which can be challenging to assess, *discover* ranks the explored subspaces and returns the top *s* solutions (i.e. top 10 in our experiments).

When applied to transcriptomic data characterizing a set of patients, *discover* identifies, for example, the genes best separating the samples, some corresponding to disease subtypes and others to various genetic similarities between patients. On scRNA-seq data, *discover* identifies the genes differentiating several cellular subtypes. Unlike most traditional clustering tools returning one prediction for the entire dataset, *discover* takes the bioinformatic analysis one step further by providing the salient genes behind cluster predictions. This constitutes a computational tool to study datasets from multiple perspectives, corresponding to the discovered subspaces.

As summarized in Figure 2, *discover* is a hybrid algorithm, combining elements from multiple evolutionary search techniques in a two-phase process. First, starting from a set of random feature subspaces, a genetic algorithm (panel a) produces new subspaces by applying feature mutations on the current set (i.e. generation) of subspaces. Next, each generation is evaluated by clustering its subspaces with a predefined algorithm and computing the internal score (i.e. Silhouette or Ratkowski Lance scores) of the resulting partition. While any clustering algorithm could be employed, we selected GMM and HDBSCAN to represent the methods requiring inputting the number of clusters in the dataset (annotated with * in experimental results) and those inferring it based on sample density, respectively. After several iterations, the best subspace is selected and passed to a maximization process (panel b). The maximization process performs a broad feature exploration and adds to the selected subspace all features that increase its internal clustering score, thus producing a larger feature subspace containing better-defined sample clusters. Performed *s* times, this process returns the best feature subspaces with their underlying clusters. Both phases (panel a, b) employ a custom feature sampling technique (panel c), involving several inexpensive statistical tests to select the most promising features preferentially, thus providing computational efficiency when the dimensionality of the input dataset is high. Further details on *discover* are provided in the annexes. Next, we present the clustering scores representing the objective function of our method.

**Fig. 2.**
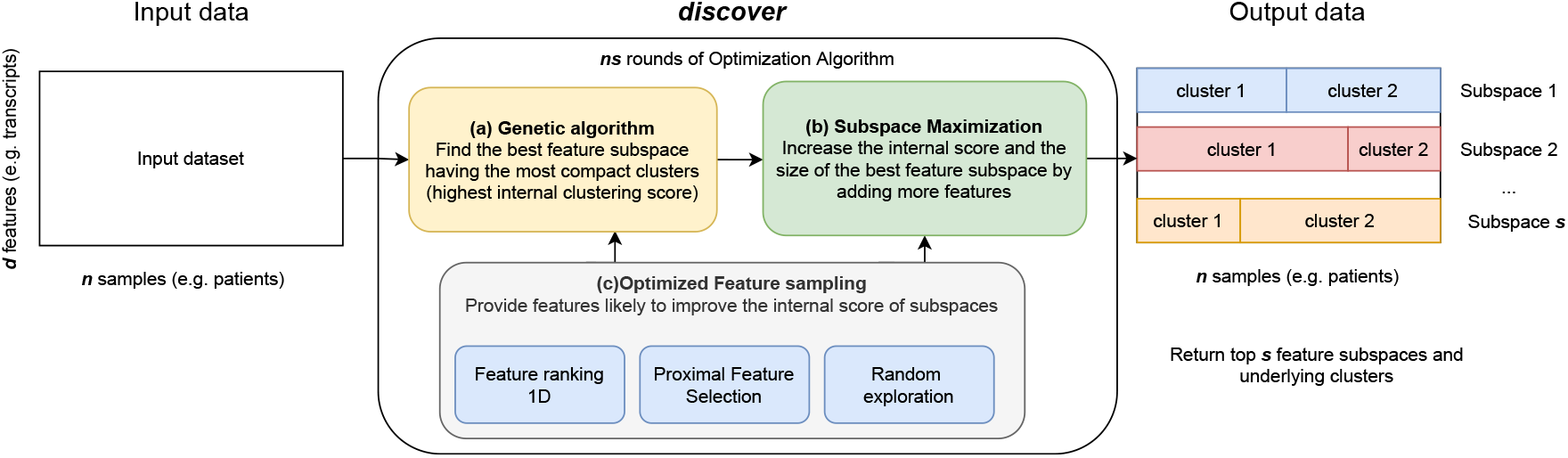
Method Overview. Our optimization algorithm returns top *s* most compact feature subspaces and the associated sample clusters (output data panel). The quality of subspaces is measured with the internal score (e.g. Silhouette scores) obtained after clustering the subspace with a predefined algorithm (e.g. GMM, HDBSCAN). The optimization algorithm runs in 2 steps. First, a genetic algorithm (box a) produces a set (i.e. a generation) of subspaces using mutagenesis and maximizes their internal clustering score. Secondly, the best subspace undergoes a maximization process, consisting of an extensive feature exploration to increase the size of the selected subspace and identify all underlying features (box b). For efficiency, both steps leverage an optimized feature selection process (box c). This process is repeated *s* times to produce the desired number of subspaces.

### 2.1 Clustering scores

Clustering analysis is typically performed when no class membership annotations (i.e. ground truth labels) are available. There are two categories of measures for assessing the performance of clustering algorithms: external and internal scores. External scores compare the predicted clustering with a ground truth annotation, provided only for validation. In contrast, internal scores estimate various properties of the predicted clustering (i.e. compactness) in the absence of external labels. Various clustering methods may create different data projections, each with well-defined clusters, and still, the resulting partitioning can be significantly different from one another [30]. To converge to a unified way of comparing clustering methods, datasets having ground truth annotations are employed for validation purposes. Most publications report external measures such as ARI (Adjusted Rand Index) or NMI (Normalized Mutual Information) scores, while the internal quality of clusters is typically presented using 2D visualizations.

This work starts from two premises: first, feature (i.e. gene) subspaces containing well-defined sample clusters may have an underlying (biological) meaning and having the computational tools to extract such subspace partitions is valuable. This assumption is also made by all clustering methods maximizing the internal quality of clusters. Secondly, the datasets may contain more than one relevant partitions and extracting multiple well-defined subspaces is valuable. Transcriptomic datasets are usually annotated for a single clustering solution. However, the samples may be grouped in multiple alternative ways (corresponding to gender, race or other genetic traits) provided the subspaces contain compact clusters. Our method identifies relevant feature subspaces by maximizing their internal score obtained by clustering each subspace. However, as demonstrated by our experiments, there is no guarantee that the most compact subspace corresponds to the ground truth. The position of this subspace in our results depends not only on the phenomena associated with the annotation but also on the impact of other complementary factors. For instance, for some conditions, race or gender could provide a better separation between patients than the disease subtypes. In this case, the first returned subspace will not be the one descriptive of the disease. The following section presents the validation protocol proposed to mitigate this problem and assess the experimental results’ quality.

## 3 RESULTS

*discover* is implemented in Python 3 and employs the feature ranking and clustering methods (i.e. GMM) provided in the scikit-learn ^1^ package. The internal clustering scores use the OpenEnsembles ^2^ package and the HDBSCAN algorithm the python implementation ^3^. For completeness, our empirical study compares the performance of experiments using GMM clustering with that of HDBSCAN. The internal clustering scores maximized by our method are Silhouette and Ratkowski Lance scores, but also two proposed adaptations, denoted as the “penalized Silhouette score” and the “penalized Ratkowski Lance score”, detailed in section A.9. These adaptations reward longer feature subspaces by adding to the original score a multiplicative factor proportional to the subspace size, encouraging the optimization algorithm to discover longer (complete) subspaces. Our method starts by pre-computing the feature ranking needed for feature sampling. Next, the optimization algorithm runs for 10 iterations and thus, returns the top 10 subspaces and their clusters. Our experiments were executed on a core i7 CPU with 16GB RAM and the underlying code is available on GitHub.

### 3.1 Validation protocol

*Validation levels*. As our method performs subspace clustering on omic data, the validation of experimental results could be performed on three levels:

1. Sample-wise, the predicted clusters are compared with dataset annotations using external clustering scores (i.e. ARI, NMI).
2. Feature-wise, the identified subspace features are compared with annotated features (if available).
3. The biological relevance of identified subspaces and clusters is assessed if underlying metadata is provided.

Performing a complete validation on a typical omics dataset remains challenging due to the general absence of datasets annotated on multiple relevant subspaces. Moreover, when using datasets labelled on a single target, there is no guarantee that the corresponding subspace appears in top s. As a solution for complete method validation, we propose a controlled data generation strategy.

#### Synthetic data

We generated datasets containing an arbitrary number of subspaces while controlling for relevant statistical properties: the subspace features, number of clusters, underlying cluster data distribution. Synthetic data makes possible the complete validation sample-wise and feature-wise. Our results are matched with the known subspaces embedded in the dataset by evaluating the feature and sample clusters overlap. The external clustering scores (i.e. ARI, NMI) are computed against the known subspace annotations for each subspace. Finally, these results are aggregated by computing an average ARI score per dataset.

#### Biological datasets

Omic datasets generally contain only one annotation allowing us to compute an external evaluation. Nevertheless, there is no guarantee of the position or presence of the target subspace in the top 10 results returned by our method. As such, the ARI score reported by our method is matched to the subspace closest to the ground truth (i.e. having the max ARI score). Because omic datasets do not annotate the most relevant features to the underlying cluster assignment, the ratio of identified feature subspaces cannot be evaluated. Thus, the method assessment is limited to the sample clustering precision, measured as an ARI score.

#### Validation with supervised feature selection

Because feature-wise annotations are missing on biological datasets, the sample labels can be used to identify a subspace associated to the ground truth by using supervised feature selection. This subspace is clustered with the same algorithms (i.e. GMM, HDBSCAN) used by *discover* to provide a comparable upper bound for our subspace identification performance (this information is not available during clustering). Supervised feature selection is also challenging as there is no consensus on the best technique [13, 40]. Furthermore, most methods require knowledge about the optimal number of features to select, which is not available. For this reason, two feature selection methods are implemented: XGBoost [8] and the mutual information score [35]. XGBoost classifies the input dataset and the feature importance is used to select the associated subspace. Mutual information measures the dependency between the target and other dataset features to capture statistical dependencies. The optimal number of selected features is determined using a grid search (50 steps) while selecting the solutions maximizing the ARI score of the underlying subspace. The supervised feature selection results are presented in the last four rows of each results table and are highlighted in gray to emphasize that this information would typically not be available.

#### Biological interpretation

The selected bulk RNA-seq datasets are accompanied by a comprehensive report of patient metadata, allowing us to establish relations between the identified subspaces and various sample characteristics, thus providing a biological interpretation to our results.

#### Competing methods

Our method is compared with relevant competing techniques for each type of dataset, also evaluated using the external ARI score. Next, we describe the four types of datasets on which our method is tested.

### 3.2 Simulated data analysis

Our data generation strategy consists in creating several feature subspaces and combining them with unrelated (i.e. noisy) features. The embedded subspaces correspond to a constructed solution to discover and have been created as multi-variate Gaussian blobs with an arbitrary number of features, clusters and co-variances while the noisy dimensions are sampled from a diverse set of distributions. Thus, we can test the validity of subspace features and associated clusters. Section A.5 details our data simulation strategy. A set of 9 datasets are generated having subspaces with 3, 6 or 12 clusters and various levels of cluster compactness. Each dataset is analyzed with 8 configurations of discover, consisting of applying GMM and HDBSCAN clustering algorithms and measuring the internal scores Silhouette, Ratkowski Lance, and their corresponding penalized adaptations. The clustering performance is assessed with ARI and NMI scores. The results in Figure 8 suggest that the penalized scores outperform the original internal evaluators on the percentage of identified features and ARI score. The penalized Silhouette score performed best when using HDBSCAN while the penalized Ratkowski Lance suited GMM best. On the simulated datasets, GMM outperformed HDBSCAN and both penalized scores providing perfect results in terms of ARI scores. For both clustering algorithms, the feature identification degrades as the number of clusters to identify and their compactness increases.

GMM (using the correct number of clusters) produces generally higher external scores than HDBSCAN, inferring this property from the sample density. HDBSCAN overestimates the number of clusters in the data, which degrades the external score. The penalized scores provide a significant performance improvement, both on ARI scores and the number of identified subspace features. For simplicity, the following experiments use only this best performing setting (GMM with Penalized Ratkowski Lance score and HDBSCAN with Penalized Silhouette).

### 3.3 Microarray data analysis

*discover* is benchmarked on 10 datasets from the Ramhiser microarray compilation, presented in section A.6. In addition, 8 competing clustering techniques, traditionally employed on microarray datasets [45] are analyzed. Affinity Propagation [20], KMeans, GMM, HDBSCAN analyze the both the entire datasets and the first 50 principal components (i.e. PCA50 + KMeans, PCA50 + GMM, PCA50 + HDBSCAN). The results in Figure 3 suggest that *discover* compares favourably with competing methods, producing average ARI scores of 0.52 and 0.27. The methods using the number of clusters as input outperform the density-based methods. As on simulated data, HDBSCAN overestimates the number of clusters in the data, degrading the external score. Performing a dimensionality reduction step before clustering does not provide a significant performance improvement. The subspaces matched with the ground truth are not the most compact on all datasets, as indicated by their ranking on Silhouette scores and Figure 3e. Additionally, the most compact subspaces (highest Silhouette scores) have generally low ARI scores, close to 0 ( Figure 3 d). A more detailed analysis is provided in section A.6.

**Fig. 3.**
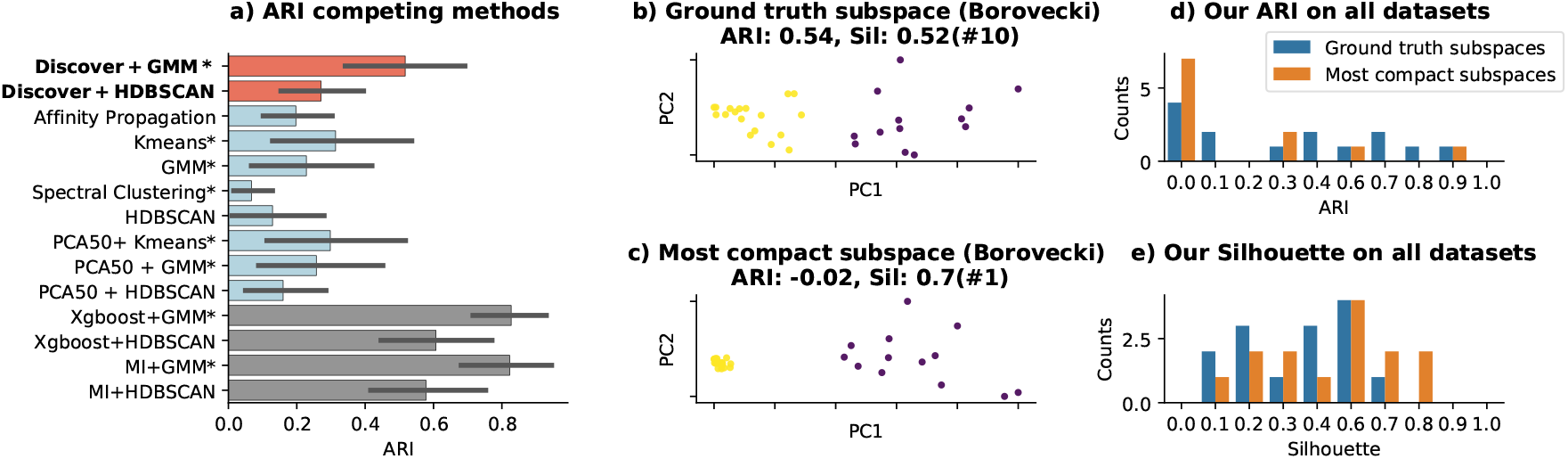
Results on microarray datasets. Our methods are compared with competing techniques on all datasets in panel a. The methods using the number of clusters are annotated by *. The bottom 4 bars are the results of clustering done using supervised features selection. Panels b, c depict 2D PCA of subspaces found by *discover*+GMM on Borovecki dataset (complete analysis in Figure 9): the subspace found to have the highest ARI wrt ground truth in panel b and the one with the highest Silhouette score in panel c. Each plot is annotated with the corresponding ARI, Silhouette scores, and the subspace’s rank by the Silhouette score (i.e. # rank position). Panel d depicts a histogram of the ARI scores for the ground truth subspace of all datasets (in blue) and the most compact ones (in orange, having the highest Silhouette). In contrast, panel e offers a similar visualization of underlying Silhouette scores.

### 3.4 Bulk RNA-seq data analysis

We analyzed bulk RNA-seq data from the Cancer Genome Atlas Program ^4^ aiming to identify cancer subtypes. Two cancer datasets (BRCA, KIRP [36], detailed in section A.7) are selected, representing breast and kidney cancer. A clinical data file provides more than 100 additional patient classifications on gender, age, smoking habits, survival patterns, additional surgical interventions, etc. Our validation method compared each subspace with all annotations and the best matching results are reported in Figure 4ab. Additionally, the same competing methods used for microarray data are assessed. Figure 5c indicates a similar method behavior: the techniques using the number of clusters as input outperform the density-based algorithms. GMM clustering outperforms HDBSCAN, which can be explained by the higher cluster compactness of the datasets, causing HDBSCAN to create larger clusters, encompassing a significant part of the dataset. The subspaces matched with the ground truth ( Figure 4a, bold) are not the most compact ( Figure 4a, in blue). Additionally, on the KIRP dataset, the second subspace matches the gender separation with high precision (i.e. ARI score 0.97) and has a higher internal quality than the subspace corresponding to the ground truth. A more detailed analysis is provided in sections 4 and A.7.

**Fig. 4.**
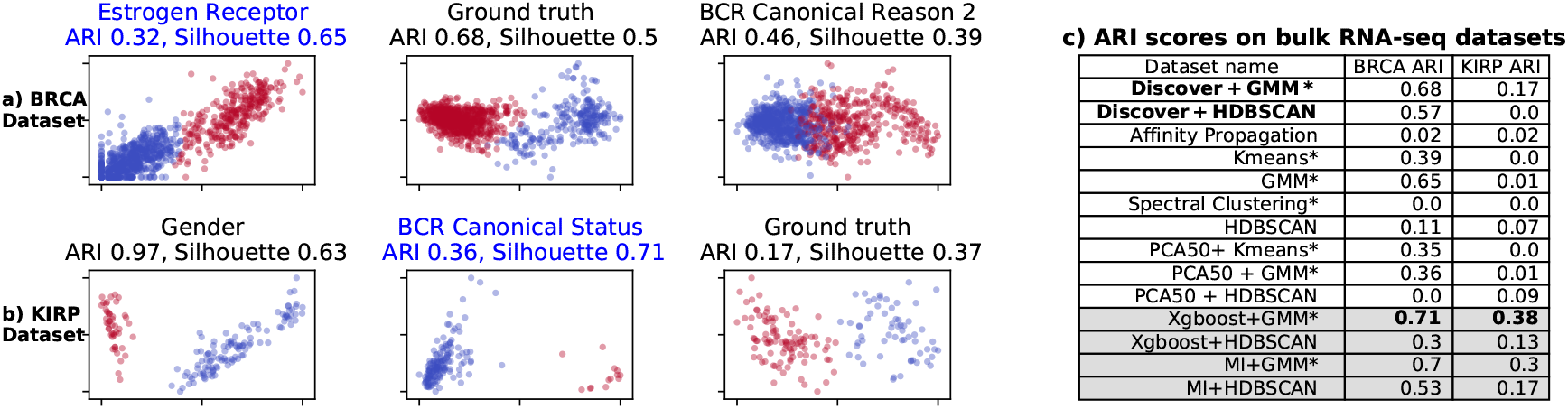
Results on bulk RNA-seq datasets (BRCA in panel a and KIRP in panel b). For each dataset, we depict the subspace aligned with ground truth, the subspace having the highest silhouete score (in blue) and a third subspace providing another example of divergent clustering. The complete analysis can be found in Figure 10. Panel c compares the results of our subspace best matching the ground truth with competing methods. The methods using the number of clusters are annotated by *.

**Fig. 5.**
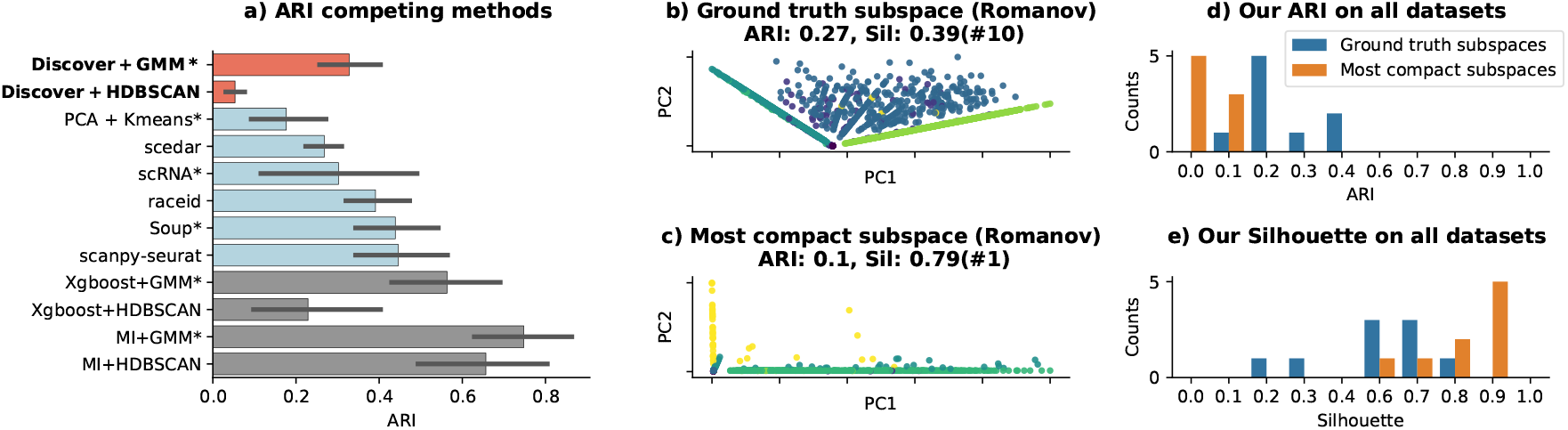
Results on scRNA-seq data. Our methods are compared with competing techniques on all datasets in panel a. The methods using the number of clusters are annotated by *. The bottom 4 bars are the results of clustering done using supervised features selection. Panels b, c depict 2D PCA of subspaces found by *discover*+GMM on Romanov dataset (complete analysis in Figure 11): the subspace found to have the highest ARI wrt ground truth in panel b and the one with the highest Silhouette score in panel c. Each plot is annotated with the corresponding ARI, Silhouette scores, and the subspace’s rank by the Silhouette score (i.e. # rank position). Panel d depicts a histogram of the ARI scores for the ground truth subspace of all datasets (in blue) and the most compact ones (in orange, having the highest Silhouette). In contrast, panel d offers a similar visualization of underlying Silhouette scores.

### 3.5 scRNA-seq data analysis

scRNA-seq data is typically affected by dropout (false zero count observations), increasing the data sparsity. Because our goal is to explore subspace search methods in a general way, we did not address the particularities of scRNA-seq data. Instead, the dropout effect is limited by filtering out all dimensions most likely affected (i.e. having most zero values).

The remaining data is still affected by dropout but to a lesser extent. Nine scRNA-seq datasets created at Stanford University from mouse cells using Smart-seq2 and 10x Genomics sequencing [50] are analyzed and detailed in section A.8. *discover* is compared with 5 competing scRNA-seq techniques: scRNA [41], SOUP [63] Seurat [49] (scanpy [58] implementation), scedar [62], raceid [42]. scRNA and SOUP require the expected number of clusters as input, the others do not.

Figure 5b reveals the lack of consensus on the best method across all datasets. Even though *discover* is not perfect and was not designed to handle the specificities of scRNA-seq data, it provides best results on two datasets. Our method performs best with GMM clustering. The ground truth subspaces are not the most compact on all datasets, as indicated by their ranking on Silhouette scores and Figure 5e. Additionally, having the highest Silhouette scores, the most compact subspaces have generally low ARI scores, close to 0 ( Figure 5d).

### 3.6 Analysis of clustering algorithms

Section A.10 evaluates the performance of several clustering algorithms as accuracy and execution time and indicates that GMM and HDBSCAN performed best on a large set of simulated datasets.

### 3.7 Analysis of internal clustering scores

Section A.9 studies several internal clustering scores when adding to compact subspace unrelated features (i.e. noisy). In short, although imperfect, internal scores can be used to identify when noisy features are incorporated into a well-defined subspace, and the proposed penalized versions of Silhouette and Ratkowski Lance score provide more accuracy for this task than the original versions.

### 3.8 Importance of individual sampling techniques

Section A.11 demonstrates that the proposed sampling methods provide a significant performance improvement compared to random exploration. The best results on sample and feature-wise accuracy are recorded when combining all proposed strategies.

### 3.9 Stability across consecutive runs

Section A.13 shows that consecutive runs of our method produce subspaces with a statistically significant feature overlap.

### 3.10 Semi-supervised subspace discovery

In addition to the presented unsupervised setting, *discover* can perform subspace clustering starting from a set of features (genes known to be associated with the researched disease) expected to be relevant for clustering. The maximization process (improving the genetic algorithm’s results) is employed to discover the subspace *S*, starting from this expected subset of its features, *S* ′, and becomes a semi-supervised problem. We compared the unsupervised and semi-supervised modes by generating 5 datasets with subspaces of various sizes (10, 30 and 60 features). The semi-supervised run starts from a random tuple of features selected from the known subspaces. The results in Figure 23 suggest that the semi-supervised mode consistently outperforms the unsupervised one for each subspace size. The biggest performance gain is achieved on smaller size subspaces, which are more challenging to identify in a high dimensional dataset.

## 4 DISCUSSION

### Validation

Our method was thoroughly evaluated (sample-wise and feature-wise) on simulated datasets, where all target samples and features are known, and our experiments reported encouraging results. However, on omic data having only one set of annotated sample labels a complete validation (feature-wise) is impossible. Therefore, a validation protocol to assess the subspace matching the provided ground truth was proposed, and a wide benchmarking exercise was performed on various types of omic data: 10 microarray datasets, 2 bulk RNA-seq and 9 scRNA-seq datasets. The ground-truth subspace was evaluated using external clustering scores (i.e. ARI, NMI). Our results indicate that *discover* compares favorably to other competing methods on all types of omic data. However, the remaining subspaces cannot be validated due to the lack of underlying feature and sample-wise annotations. Instead, we attempted to attach a biological interpretation to the discovered subspaces by leveraging the sample meta-data when available. Our analysis suggests that the remaining subspaces generally correspond to other traits (e.g. the gender subspace on KIRP). While existing bi-clustering techniques identify one cluster of co-expressed genes for all samples, the subspace clustering problem addressed by *discover* assumes that all relevant subspaces should have at least 2 compact clusters, which constitutes a different objective and limits the possibility to reuse annotated bi-clustering datasets. Thus, the partial method validation on biological data remains an open issue until subsequent annotations are available.

### Internal scores

Our empirical results indicate that the subspace associated with the ground truth is rarely the most compact one. This observation raises a problem for the clustering methods maximizing the internal quality and could explain the lack of agreement between state-of-the-art methods documented in [21]. Moreover, the most compact subspace in a dataset may correspond to a different target than what is sought after. For example, on the KIPP dataset, the discovered gender subspace has a superior internal quality than the subspace corresponding to the ground truth identifying the pathology. We hope that raising awareness to this point would encourage the creation of more datasets annotated on multiple traits and when possible, also feature-wise. This could enable the identification of what gene sets the identified clusters correspond to and in turn, would make possible a more robust evaluation of the numerous existing clustering methods.

### Biological interpretation of results

When analyzing newly sequenced datasets for which the sample annotations are missing or incomplete, the validation of computational results is challenging. To help with the interpretation of discovered subspaces, section A.12 proposes 4 strategies consisting in leveraging patient metadata, the gene ontology, databases of known genes or performing survival analysis to provide biological interpretations to discovered subspaces. This step can also provide a corrective role, which can be exemplified by analyzing the gender subspace in the KIRP dataset. Figure 21 depicts the PCA representation of these 11 features subspace and the resulting GMM clusters. We speculate that the only patient misclassified by our method (highlighted in black) has been incorrectly encoded given its distanced position to the annotated cluster. All subsets of the 11 features have been clustered, and none of them places this sample in the other cluster.

### Curse of dimensionality

As the number of features in the searched subspaces increases, our method is affected by the curse of dimensionality. A solution could be reducing the dimensionality of subspaces before clustering, when the feature size exceeds a threshold; however, it would increase the overall computational cost. Another approach is to use a clustering algorithm leveraging a distance measure suitable for processing high dimensional datasets [1], such as a fractional *L*_*k*_ norm.

## 5 CONCLUSION

The main contributions of this paper can be summarized as follows:

- Proposal of a subspace clustering method optimizing the internal clustering scores of subspaces and leveraging unsupervised feature rankings in an efficient sampling strategy. Our method can be used for unsupervised and semi-supervised problems.
- Adaptation of internal clustering scores to provide comparable results for datasets of various feature sizes.
- A broad experimental study on simulated, microarray, bulk RNA-seq and scRNA-seq data, comparing our method with competing techniques. We demonstrated that the ground truth subspace is rarely the most compact one, and other subspaces may provide biologically relevant information.

We hope that our work raised awareness of the difficulties of robust validation of clustering results and motivates more detailed annotation exercises.

## A RESEARCH METHODS

This section details our method’s components and all ablation studies performed to characterize

### A.1 Feature space sampling

Feature sampling is the functionality of providing feature candidates which, if incorporated in a given subspace *S* ′, have a higher likelihood to increase its internal score than a randomly selected feature. The proposed features are selected probabilistically from one of the three categories: (1) uni-dimensional ranked features based on statistical scores (2) the closest features to an existing subspace dimension (i.e. anchor feature) and (3) random selection from the least explored dataset features. In our experiments, an equal probability for selecting features from one of the three categories is employed. This setting combines both exploration (through random selection) and exploitation, building upon existing components in the subspace (2) or on features having relevant statistical properties. The proposed sampling technique allows our method to perform intelligent feature selections bringing computational efficiency, as relevant subspaces are discovered faster than on random exploration. Moreover, the importance of the feature selection technique increases with the size of the feature space to explore. A.11 presents an ablation study which highlights empirically the advantages brought by each component of the feature sampling heuristic and contrasts them with random exploration.

#### A.1.1 Uni-dimensional feature ranking

In this step, the input features are ranked using several uni-dimensional statistical properties, which leads to the identification of a pool of statistically important features, having a higher likelihood to contribute to well defined sample clusters. The notion of unsupervised feature importance and the underlying statistical tests have been proposed in the works of Ferreira et al. [19] and Liu et al. [38] and are detailed below. These heuristics have been combined in a three-steps process, consisting of: (i) removal of uniformly distributed features (ii) computation of spectral similarity and dispersion-related tests for each of the remaining features (iii) selection of top-scored features (here we used top 10%) across all tests with an ensemble voting strategy (i.e. we retain the features for which an arbitrary number of measures are in agreement).

As uniformly distributed features having high entropy are less likely to contribute to well-separated sample clusters, our method starts by calculating the entropy of each feature, using the relative frequencies of features’ binned values (20 bins). All features with an entropy score greater to a given threshold are tagged as non-important and will be ignored for the remaining part of the method. The second step consists of applying the statistical tests for dispersion proposed by Ferreira et al. [19] : Mean Absolute Difference, Dispersion, Mean Median and Arithmetic Mean Geometric Mean. Additionally, the spectral feature selection method described in [38] is included in our feature statistical tests, which models the global data structure using the eigen-system of normalized Laplacian matrices. Our experiments show the complementary of these methods in terms of identifying important features on both generated and omics data ( Figure 6).

**Fig. 6.**
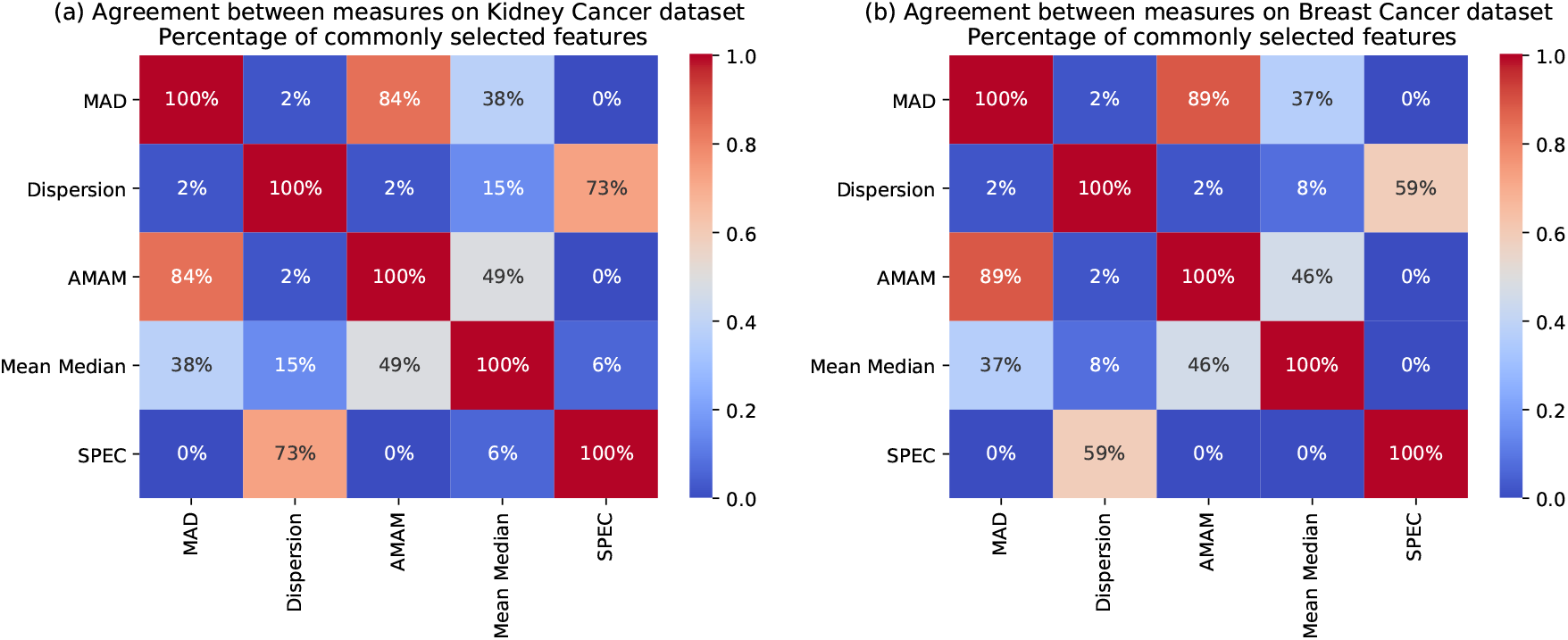
The statistical tests show complementary results in the selection of features. Here we illustrate the results on the RNA-seq KIRP and BRCA datasets and show that each measure captures diverse features. The distribution of scores is also highly dependent on the input data-set.

After each feature is evaluated with 5 statistical tests, the final step consists of translating the underlying values into a single feature ranking, providing a subset of important features. This is achieved by computing an ensemble voting for each of the tests performed in (ii): a dimension is ranked as important if it has the consensus voting of at least *m* measures, where m is an arbitrary number of statistical tests in agreement. In our experiments, we used *m* = 3 (i.e. 3 from 5 statistical tests are in agreement) but lower values for m make the method more permissive and increase the total number of returned candidates.

#### A.1.2 Sampling proximal features

Adding to a subspace *S* ′ new features which are similar (i.e. close) to its existing features is more likely to improve the compactness of existing sample clusters than incorporating random dimensions. Starting from this intuition, we first present the measure of similarity employed to quantify proximal features and then the underlying search method used to identify these close dimensions.

Omic datasets contain relevant subspaces consisting partially of co-expressed genes, having similar expression patterns. Studies like [22, 29] have shown that on gene expression data, the similarity is best captured with correlation distances. For this reason, in our experiments on omics data, we employed the Pearson correlation, but any other distance measure could be provided instead as input. Starting from the subspace *S* ′, a random feature is selected (i.e. anchor feature) and one of its k nearest neighbors [3] is proposed as a candidate for insertion. In our experiments, top 3 nearest features have been pre-computed for the entire dataset, before starting the optimization algorithm.

#### A.1.3 Random exploration

The last technique employed in our feature selection is the random exploration, which provides the possibility to integrate new features, which have not passed the statistical tests for feature importance and are not proximal to other dimensions in the subspace to optimize.

When optimizing the score of a subspace by adding new features which increase its internal clustering score, the selection of new features is done by first sampling one of the 3 presented techniques. The three feature sampling categories have equal probability to be selected. For computational efficiency, the uni-dimensional feature ranking (i) and the top 3 proximal features (ii) are computed and stored before running the proposed optimization algorithm, presented in the following section.

### A.2 Genetic algorithm

Genetic algorithms are meta-heuristics used to find optimal or near-optimal solutions to complex problems. They rely on biologically inspired operations such as mutations and crossovers in order to optimize a population of individuals and thus, identify the best offspring. Executed in an iterative way (i.e. optimizing one generation of individuals after the other), genetic algorithms allow for a vast feature exploration which makes them suitable for handling high dimensional problems.

In our method, a generation represents a set of *p* feature subspaces (*p* = 50). The initial generation of subspaces is created by generating randomly the feature subspaces and evaluating them. For efficiency, the random subspace generation gives more weight to selecting features ranked as important by the preliminary feature selection.

The evaluation of a subspace consists of clustering it with a predefined algorithm ( i.e. GMM, HDBSCAN) and computing the underlying internal quality score (i.e. Silhoutte, Ratkowski Lance). The goal of the genetic algorithm is to identify the feature subspace with the highest internal score, corresponding to the most compact feature clusters. A new generation of subspaces is produced by either combining the features of two subspaces (i.e. crossover) or by mutating the features in one subspace using feature insertions, replacements or deletions, as detailed in the following section. Performing feature mutations relies on feature sampling mechanism to provide candidate features used in the creation of new subspaces. Performing random feature sampling is an inefficient strategy when the feature space is large: a large number of operations should be performed to find the features in agreement with a given subspace. The presented feature selection mechanism makes the optimization algorithm more computationally efficient such that even a small number of iterations is able to identify solutions with good internal and external clustering scores.

The creation of the new generation produces more subspaces than the original generation (i.e. 150 new subspaces). Only top *p* (i.e. 50 subspaces) selected by their internal clustering score are kept to become parents in the next generation. The selection pressure increases with the arbitrary number of offspring to generate, from which only top subspaces become parents. Combining this low selection pressure with a high mutation rate helps maintain diversity in the population and mitigates the premature convergence [7].

In order to avoid compromising the quality of subspaces by introducing multiple noisy dimensions simultaneously, the number of features to integrate in a new subspace through insertions or replacements is limited to one at a time. This strategy slows the convergence time, as identifying a subspace S requires a number of iterations proportional to its size, which becomes computationally expensive for large subspaces. For this reason, the maximization process was introduced and allows the efficient discovery of longer subspaces. As depicted in Figure 7, this process is iterated several times (i.e. in our experiments, 3 iterations) before the best subspace is selected and passed to the next phase, the maximization process.

**Fig. 7.**
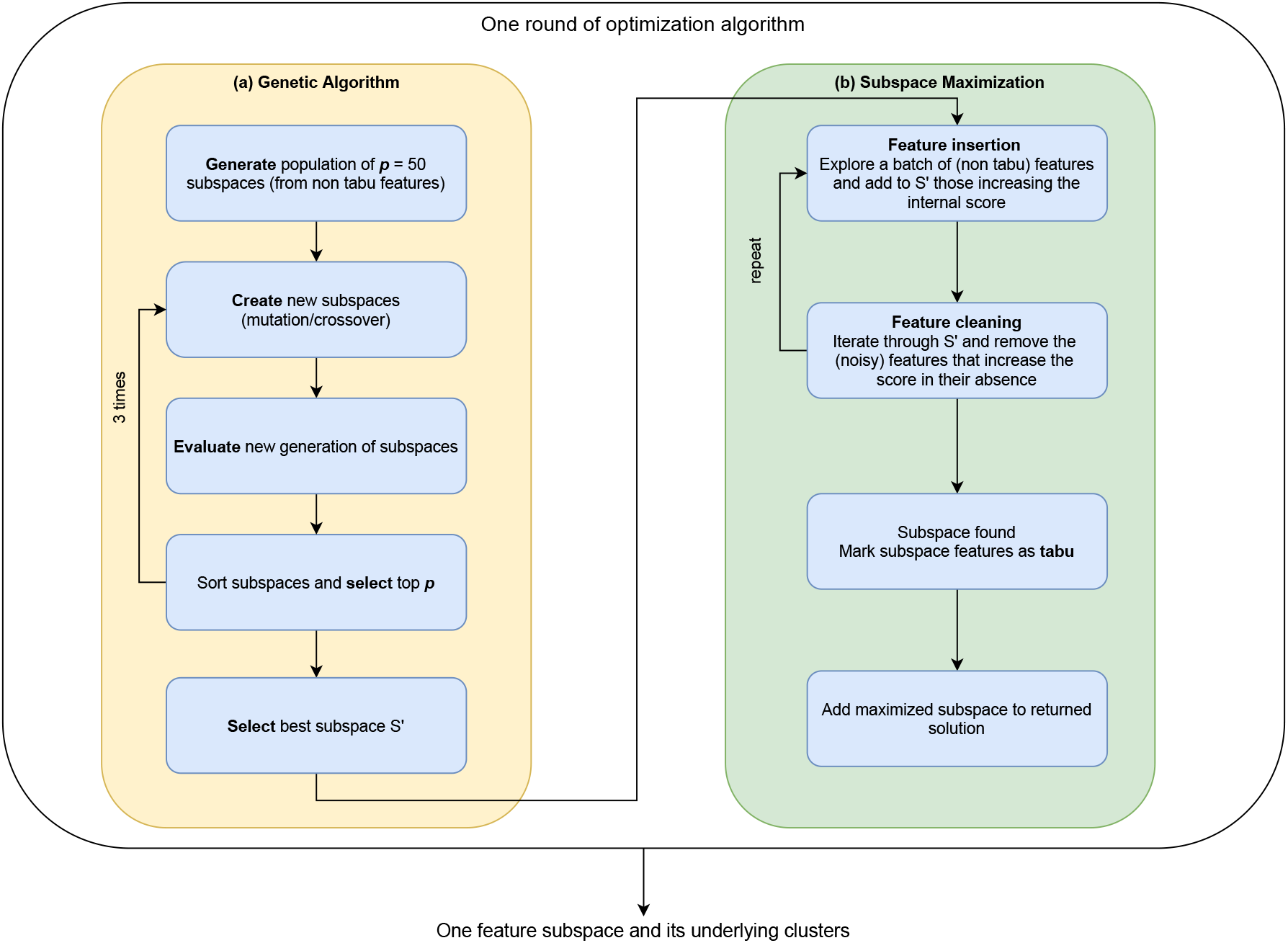
Overview of the optimization algorithm. A generation of 50 subspaces is optimized over several iterations using a genetic algorithm (panel a). The best subspace, having the highest internal clustering score, is passed to the maximization process (panel b), which consists of exploring a wide range of features and integrating those that improve the score of the selected subspace. Once the best subspace has been identified, it is added to the final solution and its features are marked as tabu (not selectable) for the next iterations. The tabu search allows the algorithm to explore and discover other subspaces with high internal scores by avoiding to fall-back on the same solution.

**Fig. 8.**
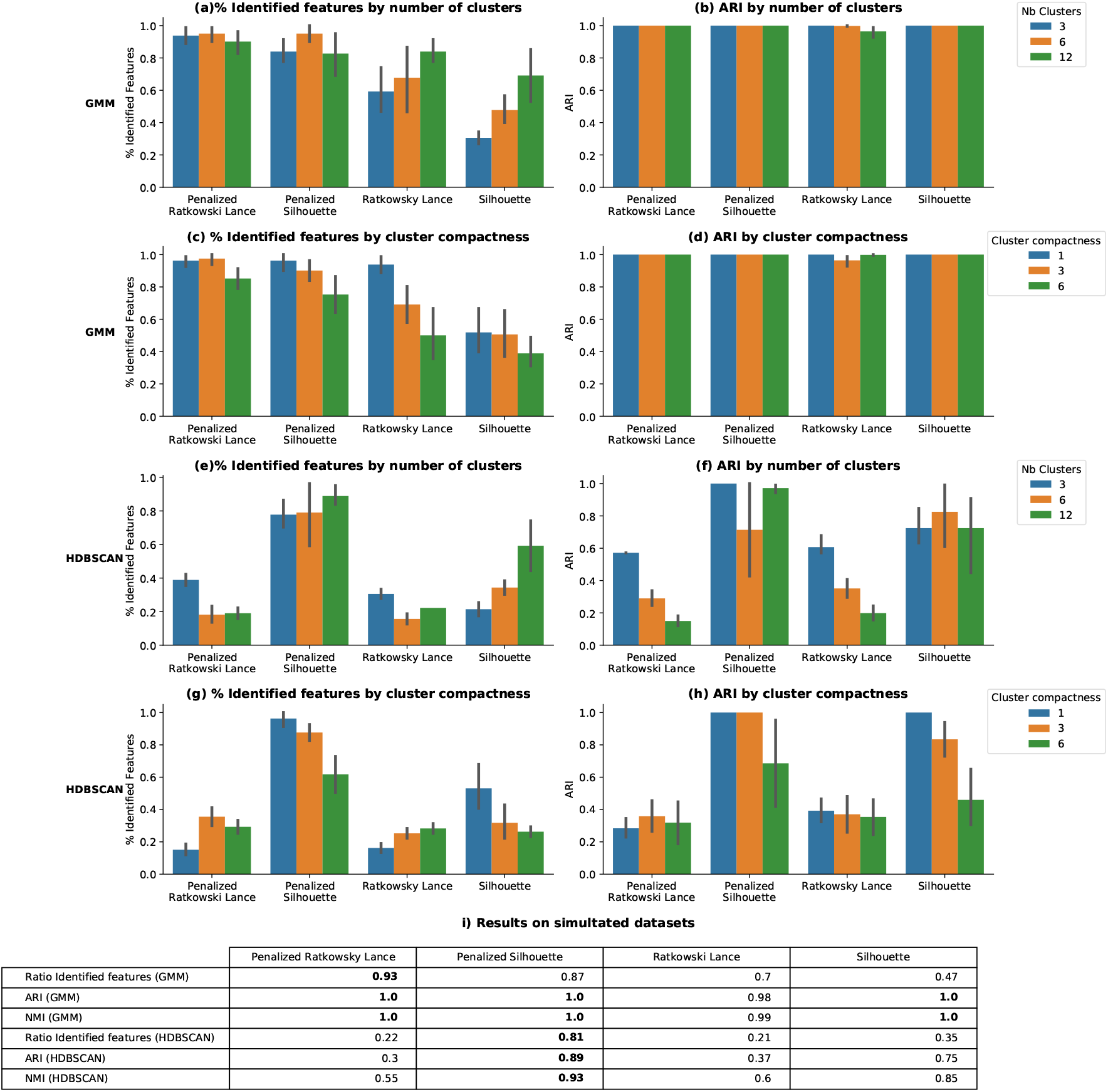
Results on simulated data, comparing 8 configurations of discover, consisting of running GMM or HDBSCAN and optimizing one of the 4 internal scores Silhouette, Ratkowski Lance and their penalized versions. 9 simulated datasets having subspaces with 3 to 12 clusters and various levels of cluster compactness (std from 1 to 6) are analyzed. The results depict the ratio of identified features (panels a, c, e, g) and the average ARI score for matching each subspace (panels b, d, f, h). The average of these results is presented in panel I, indicating that the penalized Ratkowski and GMM outperform the other configurations.

### A.3 Details on the genetic algorithm’s operations

The mutation process of a population of subspaces consists of performing an arbitrary number of crossovers, insertions, replacements and deletions operations. We start from a population of *p* subspaces, where *p* is an arbitrary user input. We perform a configurable number of each listed operations, for example *p* insertions, *p* replacements, *p* /2 crossovers and *p*/2 deletions which produce 3 * *p* offspring; after evaluation and sorting, we keep only best *p* subspaces. The configuration parameters can be adapted when having more knowledge the analyzed data-set. For instance, if it is known that the target subspaces have only a couple of features, the number of cross-over operations which allow the quick discovery of long subspaces, could be decreased in the favor of insertions and replacements. Also changing the ratio between the number of explored versus selected individuals for the next round controls the exploration rate. All operations except for the maximization process are performed on the set of non-redundant features (4).

Insertion and replacement operations incorporate one feature at a time and employ a similar process for sampling new features. The insertion process appends one feature to an existing subspace while the replacement removes one random feature and replaces it with a new one.

The new feature will be sampled with arbitrary probabilities from one of the following sources: the set of important features (1), the close features (2) or randomly (3), from the least explored non-redundant features.

The crossover operations put together the features of two randomly chosen subspaces from the selected population. Crossovers allow the rapid discovery of longer subspaces, as they incorporate multiple features at a time.

The deletion operation applied on a subspace randomly selects one feature and removes it; if the result has a higher score, it may be incorporated in the next generation and allow for more suitable features to be added. The goal of the deletion is to filter out unsuitable features from the best subspaces.

### A.4 Subspace maximization

The maximization process is an exploitation technique which attempts to completely reconstruct *S* from a subset *S* ′ by performing a large feature exploration; the dimensions which when added, improve the internal score are incorporated into the returned solution. The feature selection is supported by the sampling method described previously. An exhaustive exploration of the remaining feature space is computationally expensive for high dimensional datasets ~*d* − *len*(*S* ′) operations for testing all features). For this reason, our algorithm terminates when the internal score of *S* ′ stops increasing after maximum number of unexplored features have been analyzed (i.e. 300 feature).

The feature insertion procedure is performed greedily by incorporating immediately all candidate features which improve the subspace score. However, noisy features could also be integrated because none of the tested internal scores is perfect for identifying relevant dimensions. The cumulative effect of integrating noisy dimensions may lower the subspace score and deviate the search to a local maximum, in which case the relevant features could be ignored. The probability of integrating noisy dimensions is reduced by alternating the feature insertion of a smaller set of candidate features with corrective feature deletion (i.e. removing the incorporated features in whose absence, a higher score is achieved).

The end condition for iterations applies when no further improvement of the subspace has been achieved and the number of explored features is larger than the exploration limit described in the previous paragraph. The continuous improvement clause makes possible the discovery of long subspaces, larger than the exploration threshold (i.e. the 300 features). To prevent the algorithm from re-converging to the same subspace and to allow the exploration of lower score subspaces, the features of the solution subspaces are marked as tabu [24], (i.e. not selectable) for the scope of following rounds. Secondly, the tabu search allows the exploration in future rounds of lower score features and subspaces.

### A.5 Generated data

Using biological-only data sets for assessing the method is problematic: the sample labels are not always present and when they are, they may not be reliable and they usually correspond to one subspace; we don’t know how many other subspaces are embedded in the dataset nor what are their features or sample clusters. Biological data may be tainted with technical noise, for instance in the form of batch effect, which makes the identification of the expected subspace clusters more complicated. Cluster-size imbalance can also affect the correct identification of expected subspace clusters and is dependent on the chosen clustering algorithm. The usage of simulated datasets mitigates all these difficulties as it allows to control parameters like the number of distinct subspaces, their size, features, data distribution and the number of embedded clusters, thus providing a granular ground truth information and making the key points of our method testable.

We employed simulated data for validating our optimization algorithm and finding the optimal set of parameters. The data generation process consists of the following steps:

- we create an arbitrary number of distinct multivariate gaussian blobs subspaces *S*_*i*_ ∈ ℝ^*m*\**n*^, where *m* is the number of features in each subspace and n the number of observations in the target dataset. We randomly sample covariance matrices, which control the cluster shape and compactness. We enrich the diversity of data patterns by varying for each subspace properties such as the number of features, of embedded clusters as well as their standard deviation (controlling the compactness of clusters). The subspaces will act as the target to be discovered with the optimization algorithm. Having full knowledge about the underlying features and clusters in each subspace allows to quantify the method’s performance as both the rate of identified features and the external score of the sample clusters
- in order to simulate the high dimensionality and the difficulty to identify the target subspaces we add an arbitrary number of unrelated features. We sampled from distributions such as uniform, normal, negative binomial, gamma an arbitrary number of features, thus adding at each step a subset *A*_*i*_ ∈ ℝ^*ns*\**n*^, where *ns* is the number of added dimensions. In order to enrich the diversity of data patterns, we also sample the values of each hyper-parameter from an uniform distribution. All these features represent noisy dimensions which should be ignored by the optimization algorithm. Using this strategy of adding diverse unrelated features, we tested the performance of our method when introducing different amounts of noisy features, ranging from 100 to 30.000
- to create a more complex feature space and to get closer to the particularities of biological datasets which are rich in various types of feature dependencies, we introduce feature redundancy by creating copies of randomly selected features and adding noise. We select an arbitrary number of features nr from both important and noisy dimensions, we enrich them with noise and we obtain a subset *B*_*i*_ ∈ ℝ^*nr* \**n*^
- the dataset generated for the optimization algorithm is the reunion of all described sampling strategies *D* = *S* + *A* + *B*, *D* ∈ ℝ^*d*\**n*^, where + is the dataset union operator
- the final step normalizes feature-wise, between 0 and 1, all values in *D*

Biological datasets have specificities in terms of data distribution and feature relations. The task of closely simulating biological data of microarray or RNA-seq types can be very elaborate: [44] explained that generating realistic data requires simulating biological ground truth data, adding biological and measurement related errors and simulating the hybridization and microarray slide. [17] showed that complex models taking into account several components of the underlying physical phenomenon to simulate, can lose in flexibility and including new factors in the model becomes a difficult task. Even though generating Gaussian multivariate distributions (blobs) is easy, for more complex distributions it becomes particularly challenging. In order to better understand the complexity of biological datasets, a sub-field of research studying various types of bioinformatic data generation emerged [17, 61]. In this work we adopt a simpler solution in terms of the fidelity of reproducing the biological data, but which provides more flexibility in terms of testing capabilities and control of the generated output. Controlling the number of subspaces and their feature sizes allows to assess the subspace feature discovery rate; controlling the number of clusters and their standard deviation allows to assess the cluster identification rate and also its dependency on the compactness of clusters (given by the standard deviation); controlling the number of unrelated features added to the target subspaces allows to assess the dependency of both subspace features and clusters identification rates on the noise ratio. We also benchmark the execution time when using datasets of varying size and we can assess the relative importance of each feature sampling step to the final subspace feature and cluster identification rates. Having access to all this information is essential for the understanding and improvement of the method.

The proposed data simulation strategy is simple and makes our method testable at multiple levels; however, the most conclusive tests are on real biological input, for which we included a selection of microarray and RNA sequencing datasets, presented below.

For the experiments presented in the 3.2 section, the setup consists of generating datasets, containing 3 different subspaces of 10 features mixed with 300 unrelated noisy features. The subspaces consist of multivariate gaussian blobs and contain an arbitrary number of clusters. The generated data consists of values scaled between 0 and 1. In order to study the impact of the number of clusters in the embedded subspaces and their compactness on the performance of our method, a total of 9 datasets are generated having subspaces with 3, 6 or 12 clusters; for each setting the cluster compactness is varied over three levels corresponding to cluster standard deviation of 1, 3 and 6. Each dataset is analyzed with 8 configurations of discover, consisting of applying GMM and HDBSCAN clustering algorithms and measuring the internal scores Silhouette, Ratkowski Lance and their correponding penalized adaptations. Having knowledge about both the features and the clusters for each of the generated subspaces allows performing a complete assessment of our method. Thus, for each subspace the identification of features is assessed by computing the percentage of identified features.

### A.6 Microarray data

The data is downloadable from ^5^ but also available as an R package ^6^. This data compilation is not attached to a published paper; it has been collected from multiple studies published over the last two decades. Multiple researchers employed these datasets for benchmarking the performance of various microarray algorithms [23, 26, 43]. This data compilation is also well documented, it provides information about the target disease, references to the original studies that published the dataset and a ground truth annotation. The datasets have from 31 to 248 samples; their dimensionality varies from 456 to 22.283 features and the underlying number of clusters from 2 to 6. More details about the diseases studied by each dataset are provided in Table 1. A complete view of underlying results is provided in Figure 9.

**Table 1.** List of benchmarked microarray datasets, detailing their size, the studied disease and the reference to the issuing work

| Index | Dataset name | Sample size | Nb features | Nb clusters | Disease | Reference |
| --- | --- | --- | --- | --- | --- | --- |
| 1 | borovecki | 31 | 22.283 | 2 | Huntington's disease | [6] |
| 2 | chiaretti | 111 | 12.625 | 2 | Leukemia | [10] |
| 3 | christensen | 217 | 1.413 | 3 | N/A (Aging) | [11] |
| 4 | golub | 72 | 7.129 | 3 | Leukemia | [25] |
| 5 | gordon | 181 | 12.533 | 2 | Lung cancer | [24] |
| 6 | khan | 63 | 2.308 | 4 | SRBCT | [31] |
| 7 | sorlie | 85 | 456 | 5 | Breast cancer | [51] |
| 8 | su | 102 | 5.565 | 4 | N/A | [52] |
| 9 | yeoh | 248 | 12.625 | 6 | Leukemia | [60] |
| 10 | west | 49 | 7.129 | 2 | Breast cancer | [57] |

**Fig. 9.**
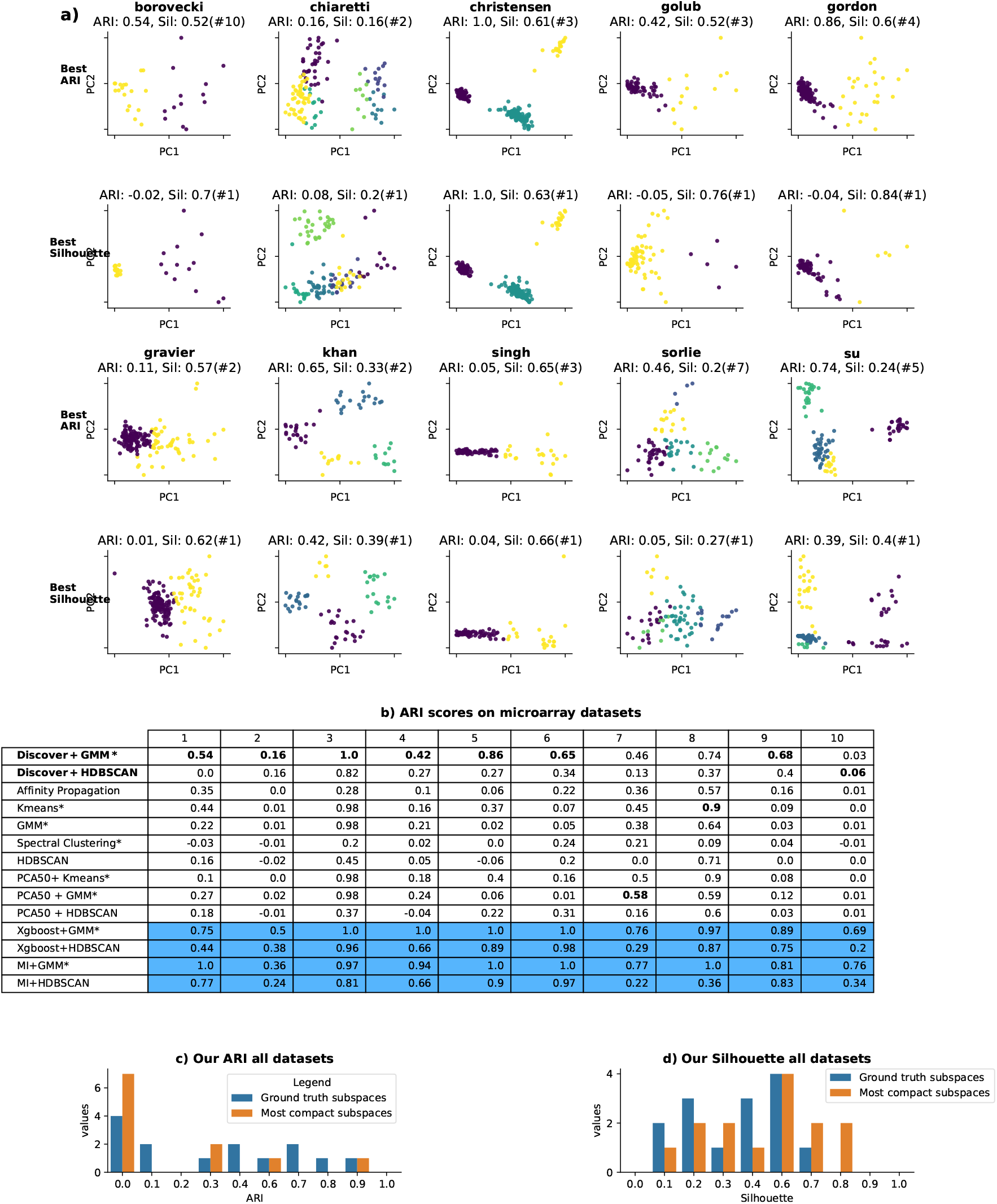
Results on microarray datasets. Panel a depicts for each dataset the subspace having the highest ARI wrt ground truth (i.e. Best ARI) and the subspace having the highest Silhouette score (i.e.Best Silhouette). Each plot is annotated with the corresponding ARI, Silhouette scores, and the subspace’s rank by the Silhouette score (i.e. # rank position). In panel b, our methods (in bold) are compared with competing techniques on all datasets. The lines in blue present the supervised features sMealencutsicorniptressuubmltsittteodgtoivAeCaMn upper bound for the expected performance. Panel c depicts a histogram of the ARI scores for the ground truth subspace (in blue) and the most compact ones (in orange, having the highest Silhouette). In contrast, panel d offers a similar visualization of underlying Silhouette scores.

### A.7 Bulk RNA-seq data

A second set of experiments targets cancer data with the goal of identifying cancer subtypes. We selected one of the best documented and widely used data compilations from The Cancer Genome Atlas Program ^7^. We analyzed two cancer datasets, corresponding to breast and kidney cancer, which were sequenced using Illumina HiSeq 2000 RNA Sequencing Version2. The datasets are accessible on the Broad institute GDAC FireBrowse Version 1.1.35 [36].They are accompanied by a detailed clinical data file which provides more than 100 supplementary type of information about each patient. Thus, we can analyze the presence of the disease (the ground truth) and other pathology specific factors, as well as patterns in gender, age, smoking habits, survival patterns, additional surgical intervations, etc. On average, half of the expected values of meta information are missing for various collection related reasons; however, we can utilize the present data to devise multiple subspace discovery validation strategies. For instance, we assess a potential correspondence between the partitions of the resulting subspaces and the clinical meta data, we perform survival analysis, gene ontology analysis and we study the presence of known cancer genes, exercises detailed in section A.12.

For both datasets we designed a common preprocessing pipeline which consists in applying a log transform, removing before the downstream analysis the genes expressed at a very low level (more than 75% of values are under 15 percentile) or have constant values and samples without subtype information. For breast cancer we also removed male samples. Further details are presented in Table 2. A complete view of underlying results is provided in Figure 10. The way in which the patient metadata was used to validate biologically our results is detailed in section A.12.1, as part of the biological interpretation subsection A.12.

**Table 2.** Overview of the RNA-seq datasets elicited for evaluating the optimization algorithm. We selected 2 data sets associated with breast and kidney cancer which have 2 embedded clusters.

| Index | Dataset name | Sample size | Number of features | Number of clusters | Disease |
| --- | --- | --- | --- | --- | --- |
| 1 | BRCA | 1213 | 20532 | 2 | Breast cancer |
| 2 | KIRP | 163 | 20532 | 2 | Kidney cancer |

**Fig. 10.**
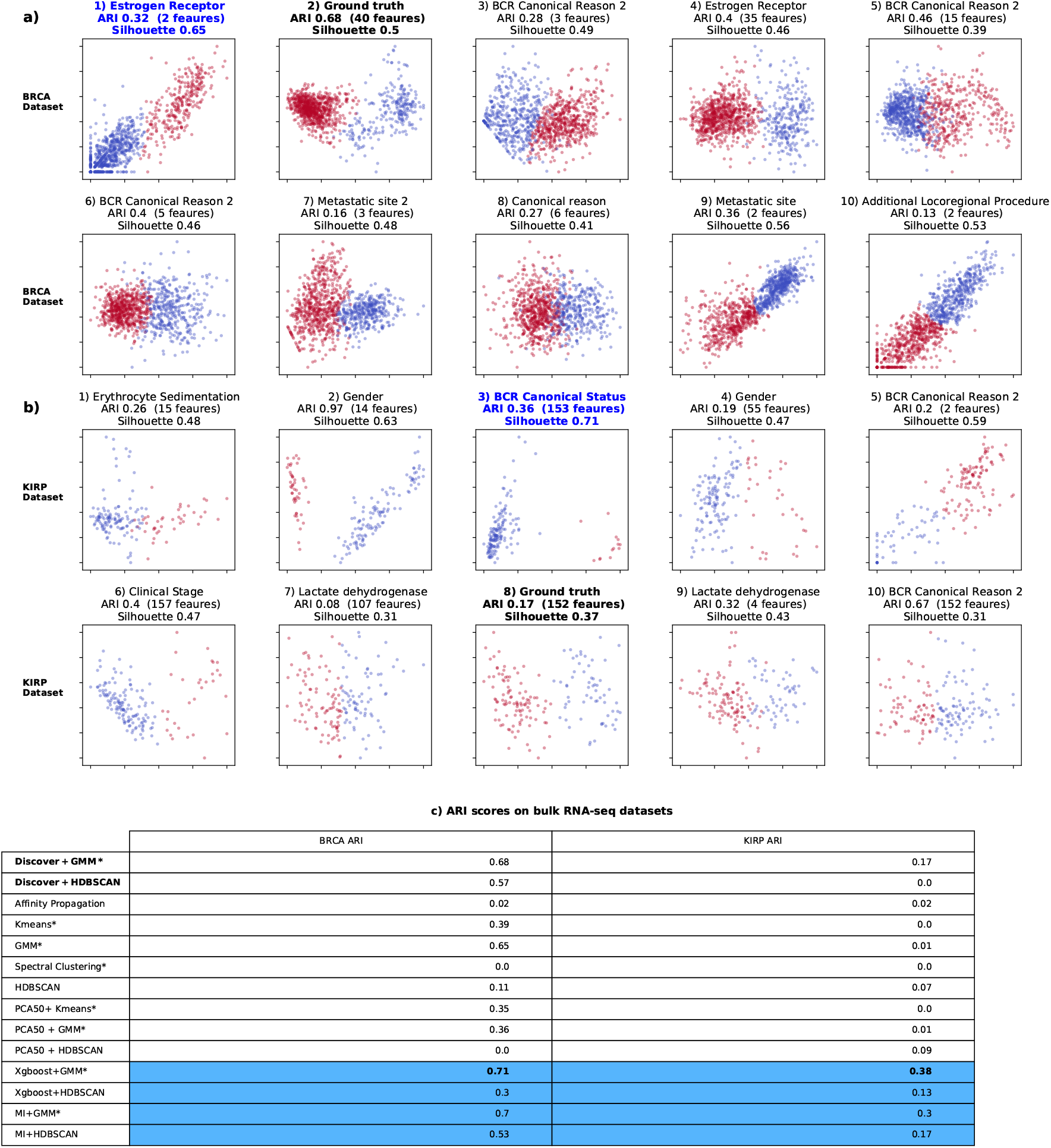
Results on bulk RNA-seq datasets. Panel a and b depict for each identified subspace in the BRCA and KIRP datasets, the best matching metadata information together with the underlying ARI and Shilhouette scores. In bold black we highlighted the subspace matching the ground truth while in blue is the most compact subspace. Panel c compares the results of our subspace best matching the ground truth with other competing methods. The lines highlighted in blue select the feature subspace in a supervised way, to give an upper bound for the expected performance of the method.

### A.8 scRNA-seq data

ScRNA-seq data is typically affected by dropout (false zero count observations), which increase data sparsity. Because the goal of this work is to explore subspace search methods in a general way, we did not elaborate on the specificities of scRNA-seq data. The dropout is addressed simply by filtering out all dimensions most likely to be affected (i.e. having most zero values). The remaining data is still affected by dropout, but to a lesser extent. 9 scRNA-seq datasets created at Stanford University from mouse cells using Smart-seq2 and 10x Genomics sequencing [50] are analyzed. The Smart-seq2 datasets have been prefixed with “Quake Smart” while the latter with “Quake10x”. Other publicly available datasets have been added, as follows: Muraro [16], Romanov [17], droplet barcoding for mouse embryonic stem cells [34], Microwell-seq for mouse bladder cells [28] ( Table 3). A complete view of underlying results is provided in Figure 11.

**Table 3.** List of benchmarked scRNA-dataset datasets, detailing their size and the number of underlying clusters.

| Index | Dataset name | Sample size | Number of features | Number of clusters |
| --- | --- | --- | --- | --- |
| 1 | Muraro | 2122 | 19,046 | 9 |
| 2 | Quake 10x Limb Muscle | 3909 | 23,341 | 6 |
| 3 | Quake Smart seq2 Diaphragm | 870 | 23,341 | 5 |
| 4 | Quake Smart seq2 Limb Muscle | 1090 | 23,341 | 6 |
| 5 | Quake Smart seq2 Lung | 1676 | 23,341 | 11 |
| 6 | Quake Smart seq2 Trachea | 1350 | 23,341 | 4 |
| 7 | Romanov | 2881 | 21,143 | 7 |
| 8 | Mouse ES cell | 2717 | 24,175 | 4 |
| 9 | Mouse Bladder cell | 2746 | 20670 | 16 |

**Fig. 11.**
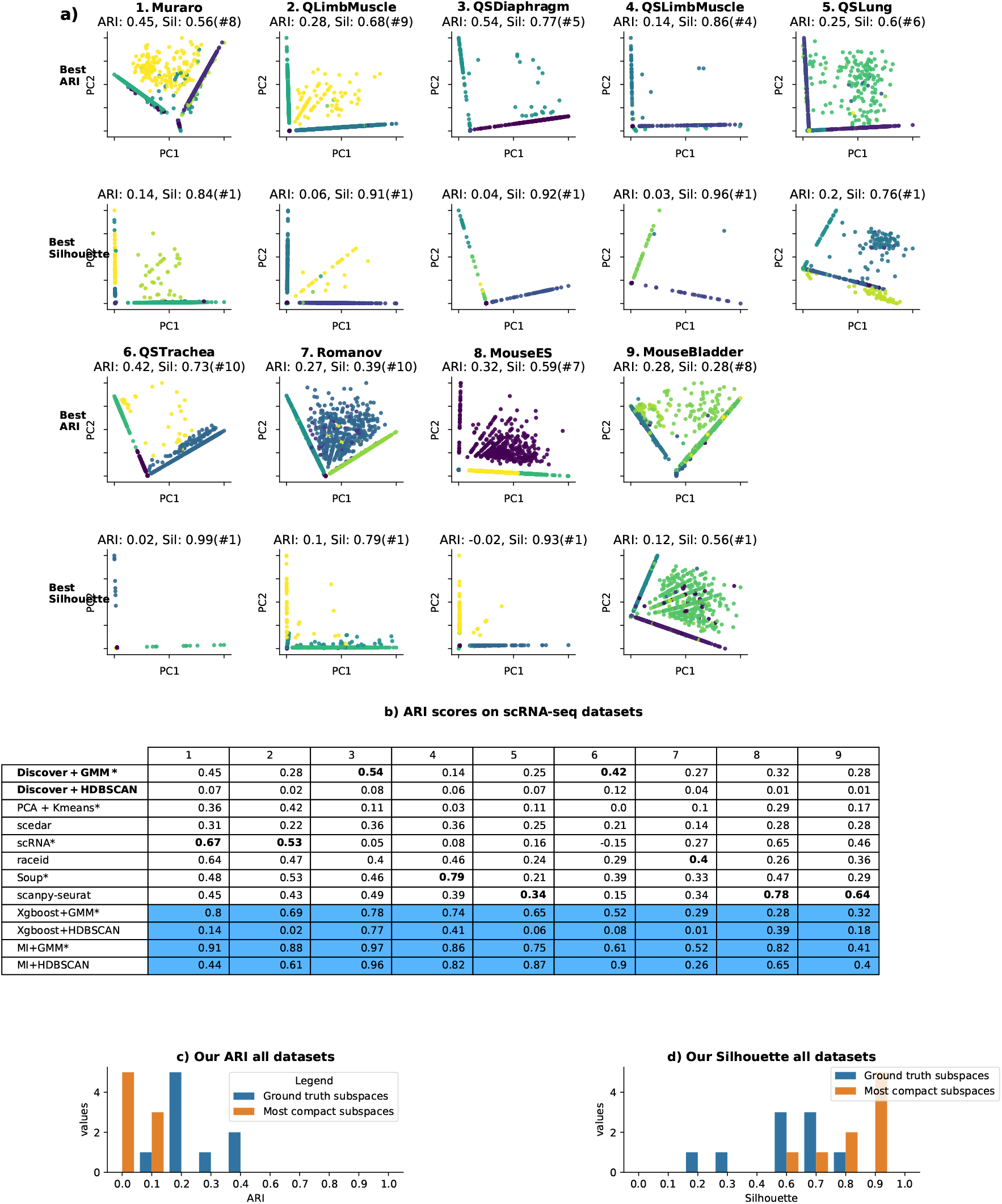
Results on scRNA-seq data Panel a depicts for each dataset the subspace having the highest ARI wrt ground truth (i.e. Best ARI) and the subspace having the highest Silhouette score (i.e.Best Silhouette). Each plot is annotated with the corresponding ARI, Silhouette scores, and the subspace’s rank by the Silhouette score (i.e. # rank position). In panel b, our methods (in bold) are compared with competing techniques on all datasets. The lines in blue present the supervised features selection results to give an upper bound for the expected performance. Panel c depicts a histogram of the ARI scores for the ground truth subspMacaenu(isncrbipltuseu)bamnidttetdhteomACosMt compact ones (in orange, having the highest Silhouette). In contrast, panel d offers a similar visualization of underlying Silhouette scores.

### A.9 Analysis of internal clustering scores

Our method identifies relevant subspaces in a given dataset by maximizing the underlying internal clustering score. This section provides a detailed analysis on the behavior of several internal clustering scores and their capacity to identify relevant feature subspaces.

An extensive report of clustering evaluators [48] is gathered in the python package OpenEnsembles ^8^. The set of over a dozen internal scores implemented in OpenEnsembles is analyzed, and for simplicity we present only the results on three representative candidates: Davies Bouldin, Silhouette and Ratkowski Lance scores. Davies-Bouldin index computes the ratio between the within-cluster distances and cluster separation. Silhouette score measures the mean intra-cluster distance and the mean nearest cluster distance for each sample. Ratkowski Lance score estimates the ratio between cluster-level dispersion and total subspace dispersion of samples in a clustering solution. The dispersion is computed as the sum of squares of the difference between the sample values and the geometric center across all features. For the last two methods, the higher the score, the better the clustering.

Next, we evaluate if internal clustering scores of feature subspaces can identify subspace clusters in a given dataset (and thus, be used as the objective function in optimization algorithms) by setting up the following experiment. A dataset has been generated with one embedded subspace *S* and additional unrelated features. Starting from subsets of *S* (i.e. random groups of features from *S*) new subspaces are created and evaluated by adding either the remaining features of *S* or unrelated features. We refer to the first category of features, which complete the original subspace *S*, as relevant and the second as noisy. The subspace evaluation consists of clustering it with a predefined algorithm (i.e. GMM) and computing the internal score of the resulting partitioning. This experiment has been performed on both simulated datasets and biological microarray data. The simulated datasets consist of multivariate gaussian blobs for which the subspace features and the unrelated noisy features are know, as they are generated following the procedure described in the data simulation section. Because omics datasets do not offer annotations for relevant features, we employed the supervised feature ranking resulting from training a XGBoost classifier to predict the ground truth annotation. Our experimental results indicate that the subspaces identified with underlying important features consistently produce the highest external scores. In this setting, the relevant features are those having the highest feature importance while the noisy ones are those having the lowest feature importance scores. Two microarray datasets (Khan, West) are selected and top 5 most important features are considered relevant while the bottom 5 are considered noisy. From all 11 studied microarray datasets, Khan and West have the highest difference between the ARI scores of clustering the relevant features subspace and the noisy feature subspace, which confirms the separation between the two groups of features. If the features considered noisy are aligned (i.e. coming from the same subspace) with the relevant ones, this experiment will fail to assess the capacity of internal scores to discriminate between features.

For each of the analyzed datasets, the internal scores of subspaces created by adding to subsets of various sizes (2-5 features) of *S* either relevant ( Figure 12, left boxplot) or noisy features ( Figure 12, right boxplot) was computed. The results indicate that the score of the new subspace decreases in over 90% of the cases when adding a noisy feature while it increases when adding a relevant feature in over 60% of cases. This experiment confirms that, even though not perfect, the studied internal scores can separate noisy features from subspace relevant features (i.e. they have a good specificity) but they may fail to identify all subspace relevant features (i.e. lower sensitivity). On average, Silhouette and Ratkowski Lance scores discriminate best between noisy and relevant dimensions; for simplicity, the rest of this analysis studies only their behavior.

**Fig. 12.**
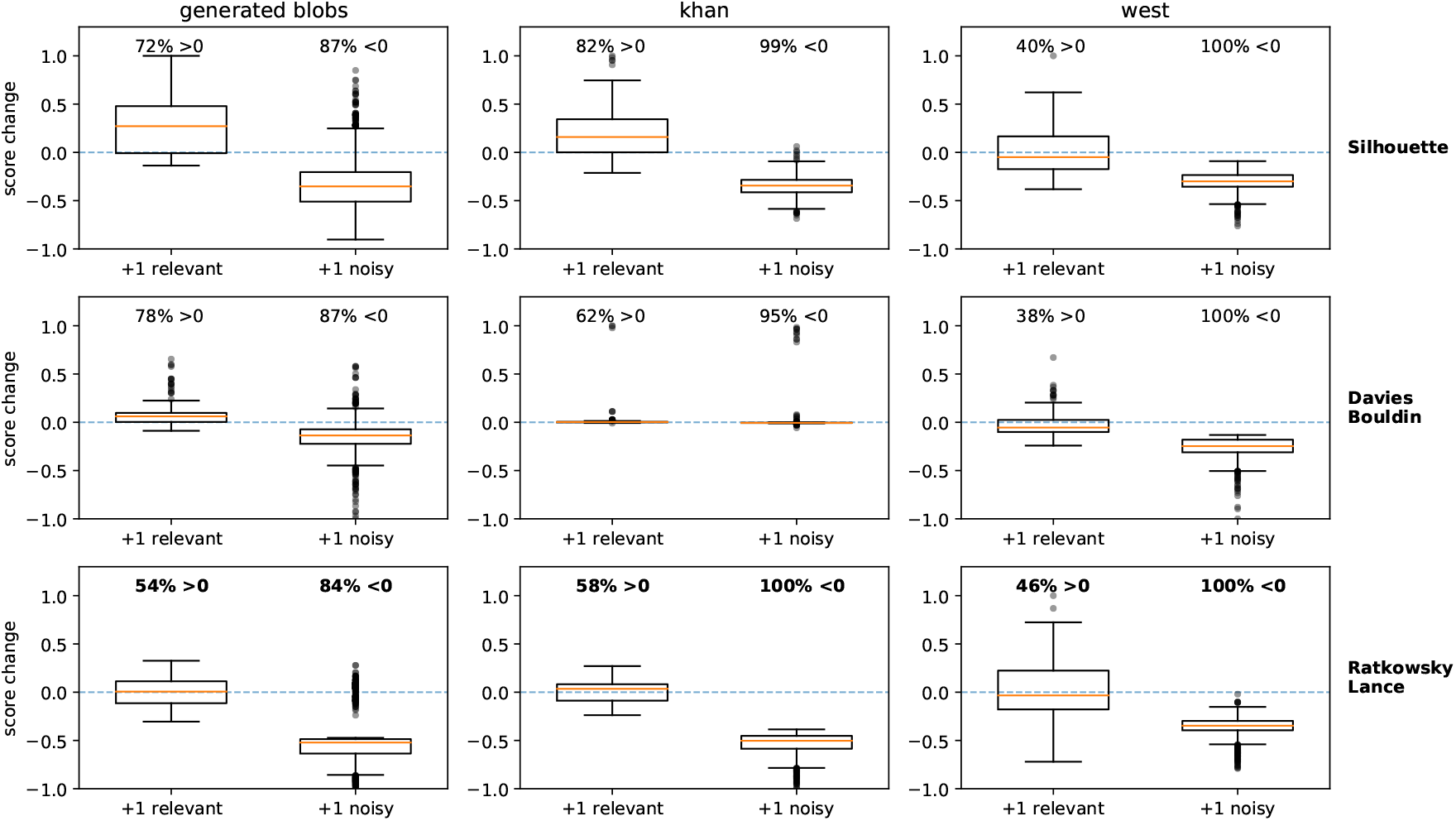
Analysis of the score change when adding relevant (left boxplot) versus noisy (right boxplot) features to a subset of predefined or top-ranked subspaces. The score change is computed when appending to subspaces of various sizes (2-5 features) one relevant or one noisy feature. The performances are evaluated on both simulated (generated blobs) and omics (Khan and West) datasets using the Silhouette, Davies Bouldin and Ratkowski Lance scores. For each experiment the percentage of relevant features that improved the score (change >0) and of noisy features that lower the score (change <0) are reported.

Next, an new experiment is devised to better diagnose the lower sensitivity of internal scores, which leads to an average of 40% of subspace features producing a lower score and prevents the full identification of the subspace features. The goal is to study if the cardinality of a subspace has an impact on its internal score. For this purpose, a larger feature subspace (40 features) is generated (i.e. consisting of multivariate gaussian blobs) and the scores of all its subsets with sizes varying from 2 to 40 are computed. Figure 13a shows that the maximum values of the Ratkowski Lance scores decrease as the subspace size increases, while Silhouette maintains relatively constant values, which explains the lower scores for the integration of relevant features, presented in the previous experiment. By using in a subspace search problem an objective function that produces lower scores when incorporating relevant features, higher scores will not necessarily correspond to the larger or more complete subspaces. Moreover, partial subspaces may have the maximum score, which creates a tendency for splitting the largest subspaces. Conversely, if the method provides higher scores when incorporating noisy features, larger subspaces will be produced, but of a lower quality.

**Fig. 13.**
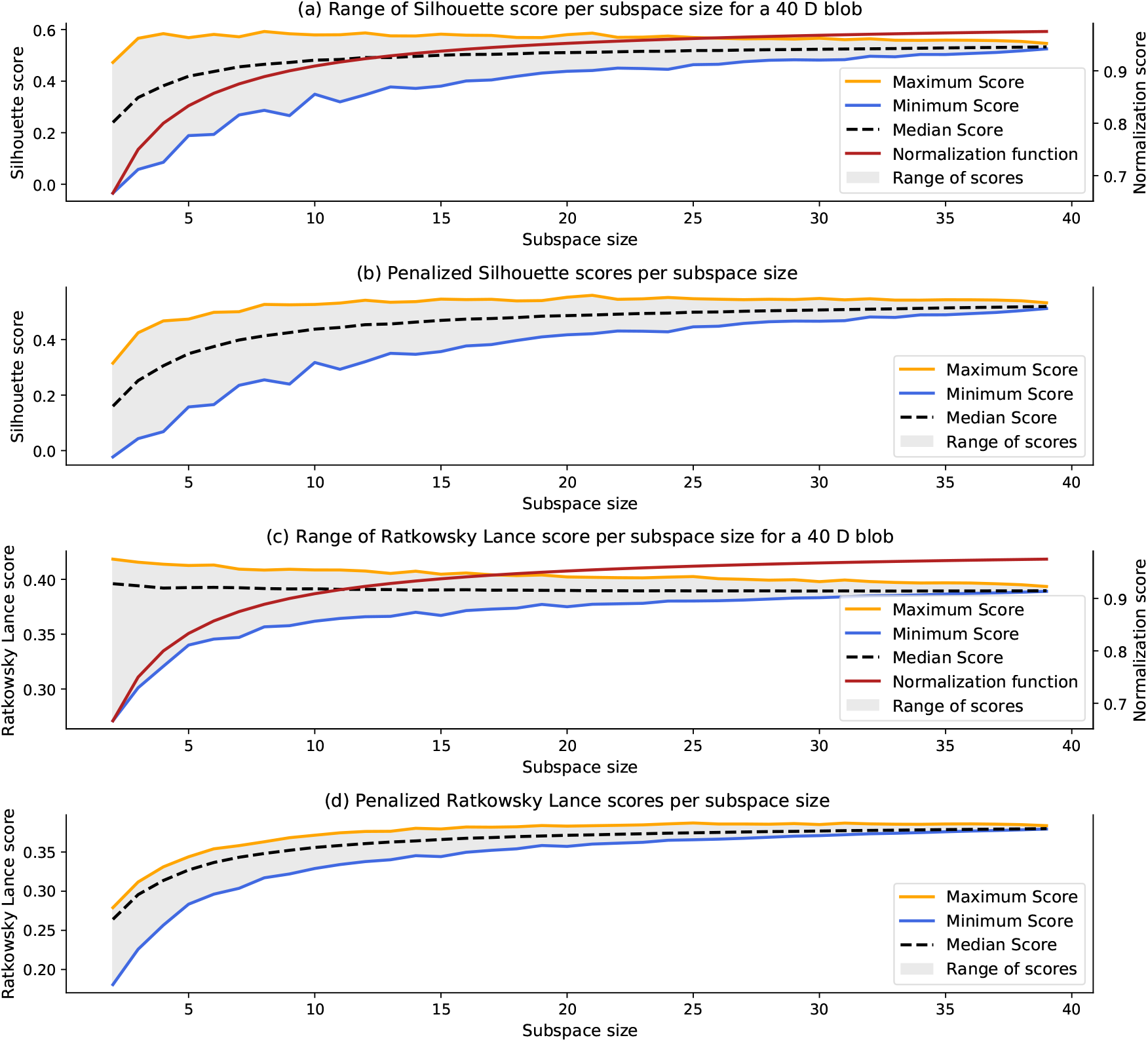
Adaptations of Silhouette (a and b) and Ratkowsky Lance (c and d) scores for penalizing small subspaces and encouraging the incorporation of additional relevant features. Ratkowski Lance displays a decreasing trend of the maximum scores (panel c, orange line) while the Silhouette score is almost flat (panel a, orange line). These characteristics limit the incorporation of additional relevant features and the discovery of large subspaces. The proposed penalization function induces an ascending maximum scores trend (panel b and d, orange line), which encourages the discovery of growing size subspaces and is labelled here “Normalization function”.

In addition to comparing the internal quality of analyzed subspaces in order to identify the most compact ones, our algorithm should also find all related subspace features. For example, if subspace S contains 3 features (i.e. *f* 1, *f* 2, *f* 3), the scoring function maximized by the optimization algorithm should return lower values for all subgroups of two features than for the entire subspace (i.e. *score* (*f* 1, *f* 2) < *score* (*f* 1, *f* 2, *f* 3)). When this condition is not met and the maximum score is attributed to a subset of the subspace, the optimization algorithm will not be able to identify S completely. In order to encourage the exploration of larger feature subspaces and allow the complete discovery of the embedded subspace, *S*, we penalized the score for smaller subspaces. A multiplicative factor *d*/(*d* + 1) is added to the original internal scores, which encourages the accumulation of features. These adaptations are denoted as the “penalized Silhouette score” and the “penalized Ratkowski Lance score” and are depicted in Figure 13 under the name “Normalization” function. The original and the penalized Ratkowski Lance and Silhouette scores are compared in Figure 14 using the same setting as in Figure 12. The results on the employed dataset show that the penalization did not compromise the method’s ability to discriminate noisy features and improved the rate of identifying essential features. The penalized Ratkowski Lance scores outperformed penalized Silhouette. The results also showed that the penalized score is not perfect (there are still noisy dimensions incorporated and relevant features ignored) but brought a significant improvement. On average, 25% more relevant features were discriminated correctly while only 2% more of noisy dimensions increased the score.

**Fig. 14.**
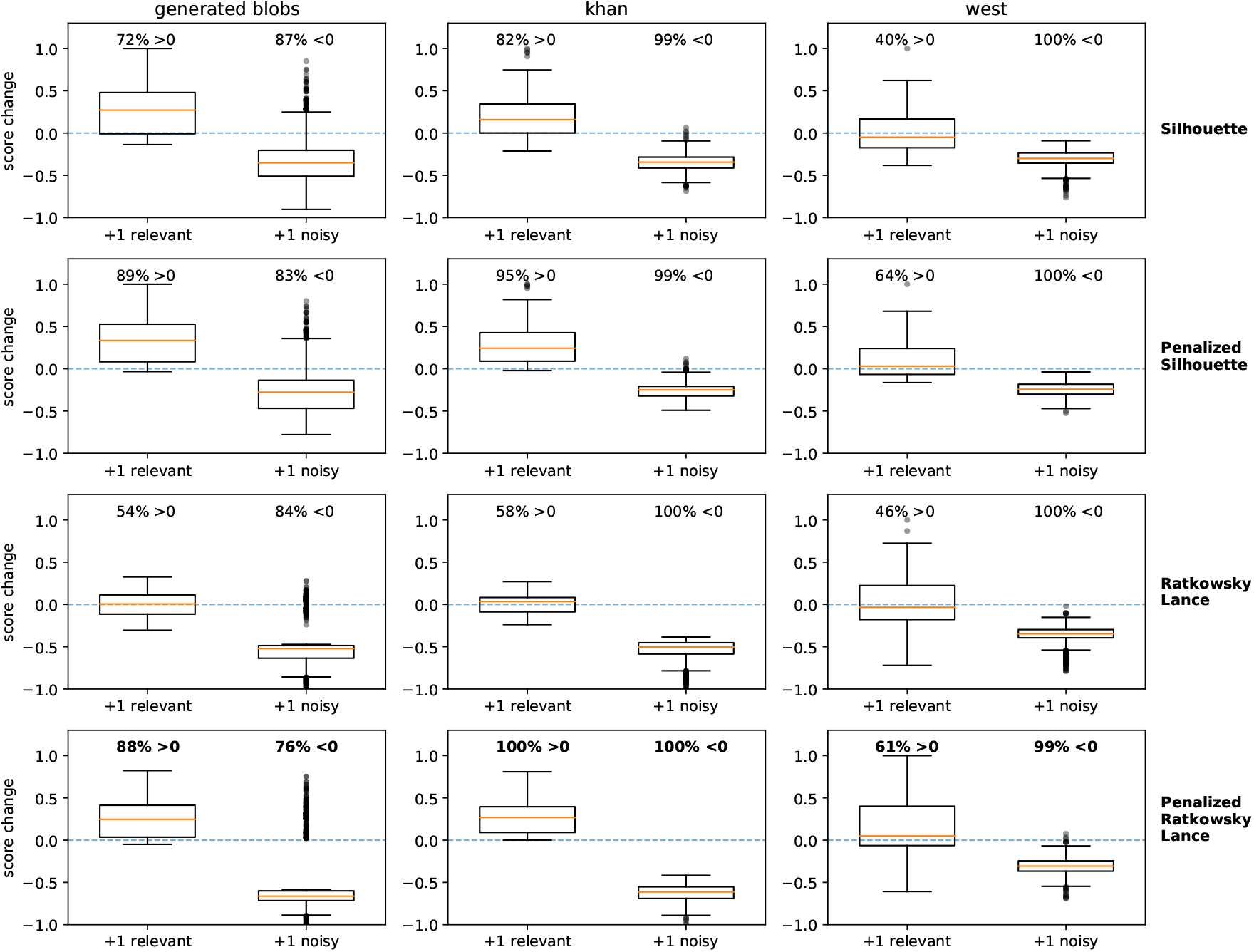
Score changes when appending one relevant or one noisy feature to subspaces of various sizes (2-5 features) using the raw or adapted scores. Original Silhouette and Ratkowski Lance scores (first and third row) are compared to the proposed penalized (second and fourth row) versions. The proposed function brings on all datasets an improvement to the possibility of discriminating, score-wise, between the addition of a relevant versus a noisy feature.

We then compared the performance of these internal scores, both in their original and penalized expressions on simulated and omics datasets, clustered with GMM and HDBSCAN. Unlike HDBSCAN, GMM expects the number of clusters as input, value which may not be known beforehand. The typical way to compute it requires specifying a set of likely values, performing a clustering for each one and using an internal quality measure to select the best partitioning. We assessed whether the selected internal scores could be used to identify the optimal number of clusters in a given subspace. We generated datasets having an arbitrary number of clusters, and computed the internal scores when clustering with GMM using the known parameter but also smaller and higher values. As depicted in Figure 15, Ratkowski Lance score tends to reward the smallest number of clusters. This shortcoming makes it unsuitable for inferring the optimal number of clusters in an unknown dataset. However, it can still be used in combination with HDBSCAN or when the number of clusters is known. Silhouette is a better candidate for this task, despite its tendency to underestimate the large values for the number of optimal clusters.

**Fig. 15.**
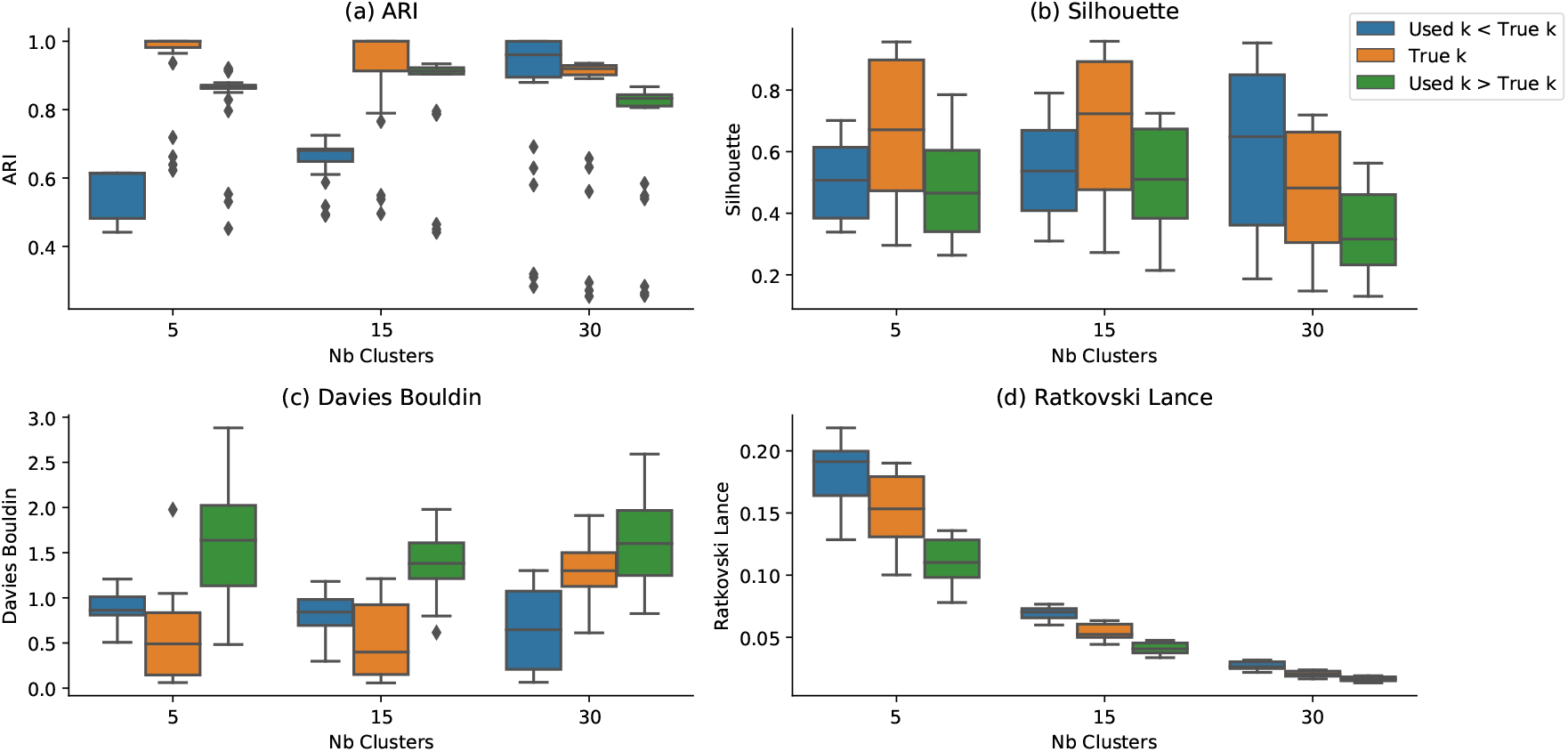
Influence of the initial number of clusters using the GMM clustering algorithm on the external and internal evaluators. We compared the external ARI (a) score with internal Silhouette (b), Davies Bouldin, (c) and Ratkowski Lance (d). The comparison is performed on datasets having 5, 15 and 30 clusters and clustered them with GMM using either the correct number of clusters (orange) or smaller (blue) or larger values (green). Ratkowski Lance always rewards the smallest number of clusters and is unsuitable for inferring the optimal value. Even though imperfect, Silhouette score is the most suitable candidate measure for this task.

### A.10 Analysis of clustering algorithms

In this section, multiple clustering algorithms are analyzed in order to identify the best performing candidates in terms of computational cost and clustering accuracy. This exercise supports the choice of algorithm to integrate in our method, where clustering is part of the subspace evaluation step. As as such, the clustering step is an operation which has to be performed a large number of times and has a strong impact on the overall method performance, both as accuracy and execution time.

Five of the most popular clustering algorithms are chosen for this exercise: Kmeans, Spectral clustering, Gaussian Mixture Models [5], Mean Shift [14] and HDBSCAN [39]. The algorithms are tested on 80 simulated datasets, consisting of gaussian blobs with an arbitrary number of samples (i.e. 450, 900, 1300 samples), of features (i.e. 3, 10, 20 features), of clusters (i.e. 5, 15, 30 clusters) and of cluster compactness (i.e. standard deviation of 0.01, 0.06, 0.12). This experimental setting allows us to quantify the impact of each controlled parameter (i.e. the number of clusters, features, samples and the cluster compactness) on the execution time and Adjusted Rand Index (ARI) score, computed with respect to the generated ground truth. The result of clustering algorithms is dependent on its configuration parameters which should be adapted to the particularities of the input data to analyze: the first group (i.e. Kmeans, Spectral clustering and Gaussian Mixture Models) require knowledge about the number of clusters to be found while the second group (i.e. Mean Shift and HDBSCAN) employ dedicated measures such as the bandwidth (MeanShift) or the minimum number of samples in a cluster (HDBSCAN). For the first group, the known number of clusters is passed the input parameter; for HDBSCAN the minimum cluster size is set at 10 and for MeanShift the bandwidth is 0.2, adapted for the input values normalized between 0 and 1. While it may be possible to obtain better results with a dedicated parameter optimization step for each dataset, this would introduce an undesirable computational overhead for our method.

A detailed analysis of the computational analysis is provided in Figure 16, depicting the dependency of the execution time for each algorithm on the number of samples, features and clusters. The execution time of the first group of clustering algorithms increases linearly with the number of clusters to be found while the density-based algorithms are less impacted. An increase in the number of samples affects similarly all algorithms. Across all experiments, the fastest algorithms are GMM (from the first category) and HDBSCAN from the density based group.

**Fig. 16.**
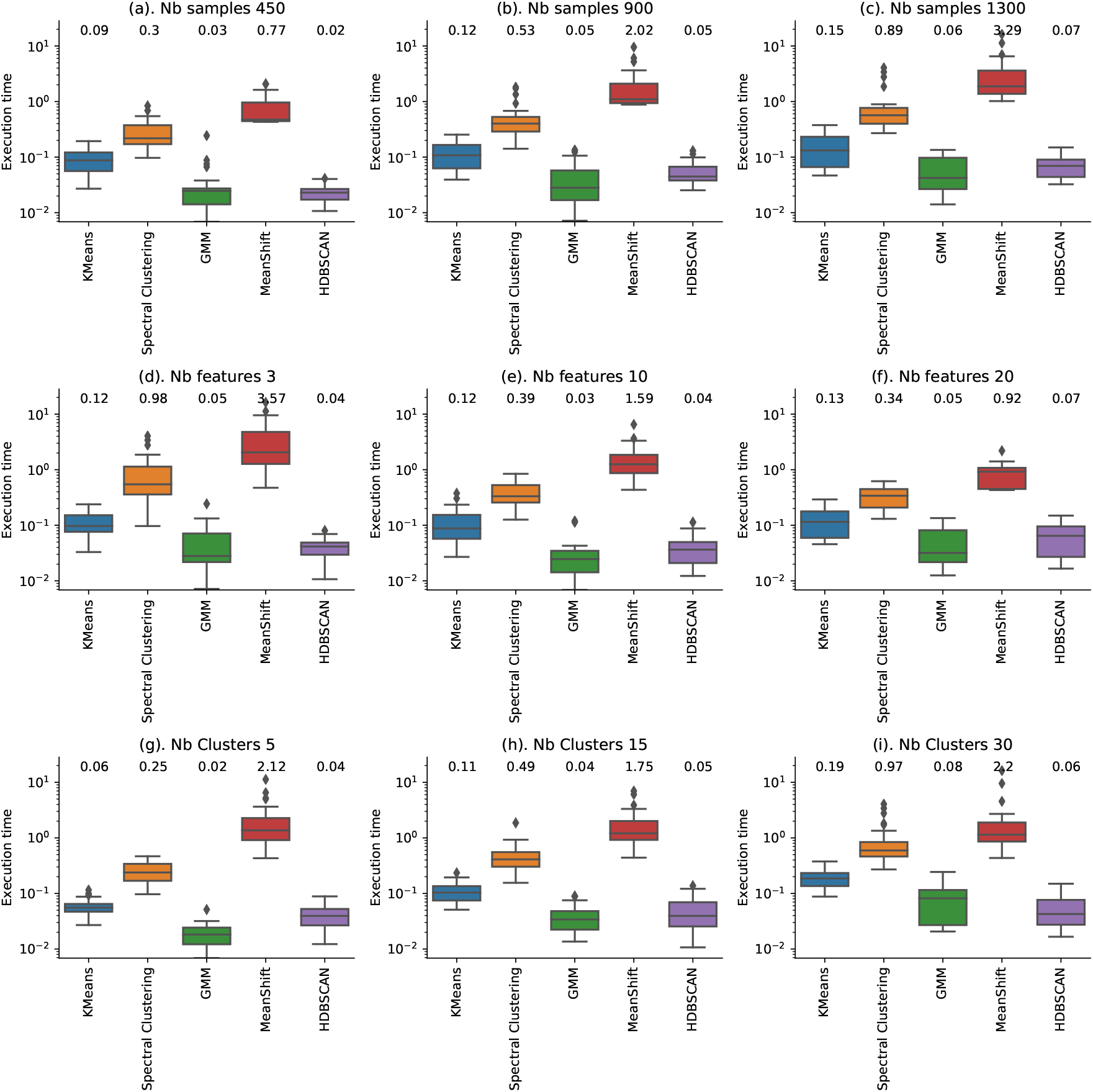
Log scale distribution of the execution time of clustering algorithms dependent on the number of samples (a, b, c), the number of subspace features (d, e, f) and the number of clusters (g, h, j). The values depicted at the top of each plot represent the average score per clustering algorithm. GMM and HDBSCAN are the fastest algorithms. GMM scales linearly with the number of clusters and the number of samples while the density based algorithms are less affected by the number of clusters.

Next, the impact of the cluster compactness (as the blob standard deviation), the number of clusters and features on the ARI score is depicted in Figure 17. As expected, the more compact the clusters the better the performance across all experiments. All algorithms perform better when the number of clusters to be found is small and the number of features in the analyzed subspace is large. However, the clustering algorithms in the first group outperform the density-based algorithms when the clusters become less compact or when the number of clusters to be found increases. Thus, if there is prior knowledge about the number of clusters in the expected subspace, it is preferable to employ one of the algorithms in the first group. If there is no knowledge about the number of clusters in the dataset, this parameter can be determined by performing the clustering several times, for a range of plausible values. The optimal number of clusters is typically identified by selecting the value corresponding to the most compact partitioning, having a maximum value for an arbitrary internal evaluator (i.e. Silhouette score). However, this procedure introduces a computational overhead proportional to the range of possible values. This overhead can be reduced by staring the analysis with a density-based algorithm in order to reduce the search space.

**Fig. 17.**
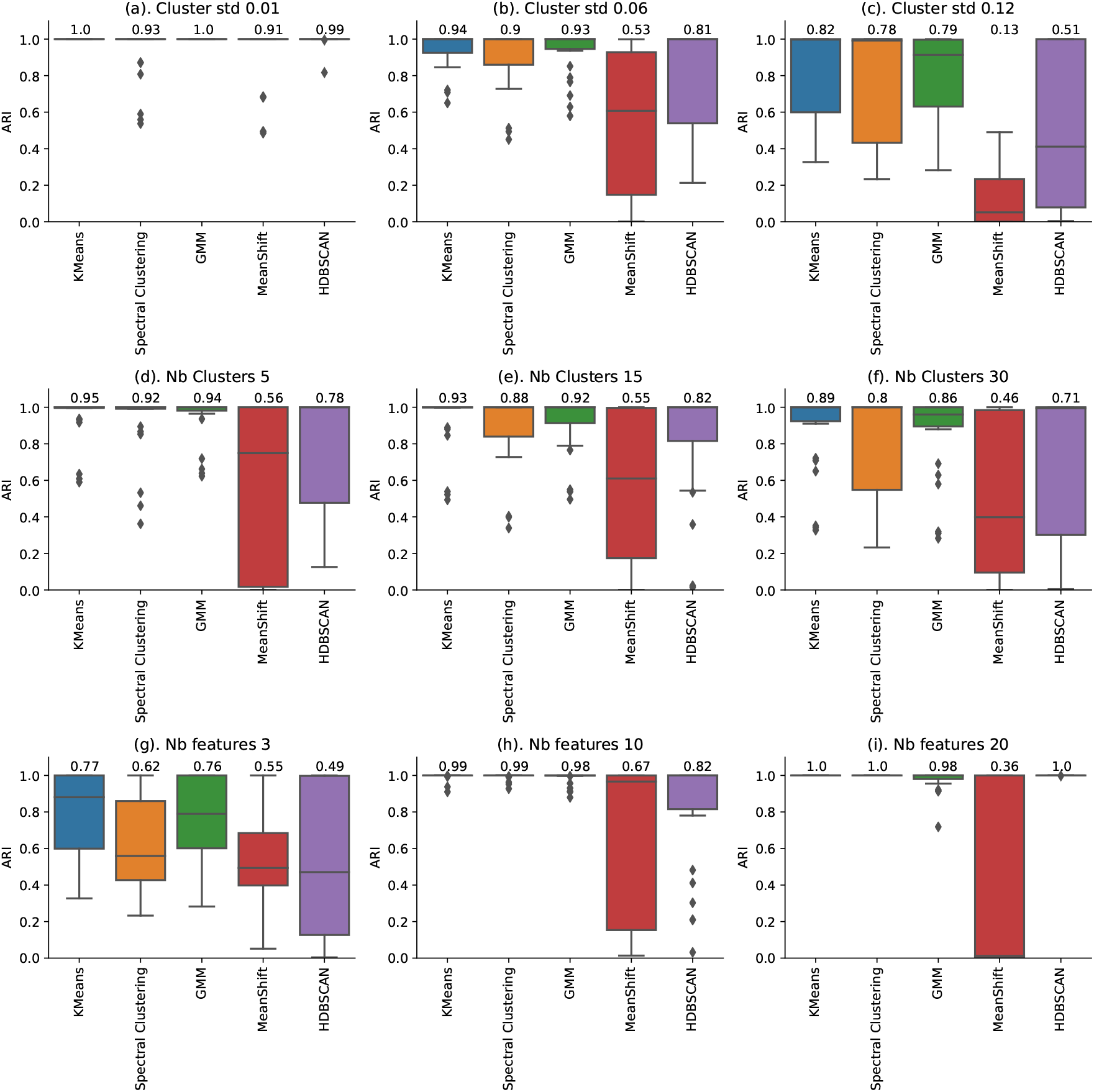
Performance of clustering algorithms measured as the adjusted rand index with respect to the ground truth, dependent on the cluster compactness (a, b, c), the number of clusters (d, e, f) and the number of features in each subspace (g, h, i). The values depicted at the top of each plot represent the average score per clustering algorithm. All algorithms perform best when the clusters are most compact. The clustering algorithms relying on the number of clusters to be retrieved (Kmeans, GMM, Spectral clustering) outperform the density based algorithms. The performance also decreases as the numbers of clusters in subspaces increases.

All experimental results are combined in an aggregated view ( Figure 18), depicting the average results across all datasets in terms of execution time (panel a) and clustering accuracy (panel b).The GMM algorithm provides ARI scores marginally lower than KMeans but bring a performance gain two times higher. HDBSCAN brings the best computational performance from the second group at a similar speed to GMM. These algorithms provide the best trade-off between efficiency and precision, justifying their choice as default parameters for all experiments presented in this work. Moreover, they represent both clustering usage settings, when the number of clusters is know and also the exploratory context, when it is inferred based on sample density.

**Fig. 18.**
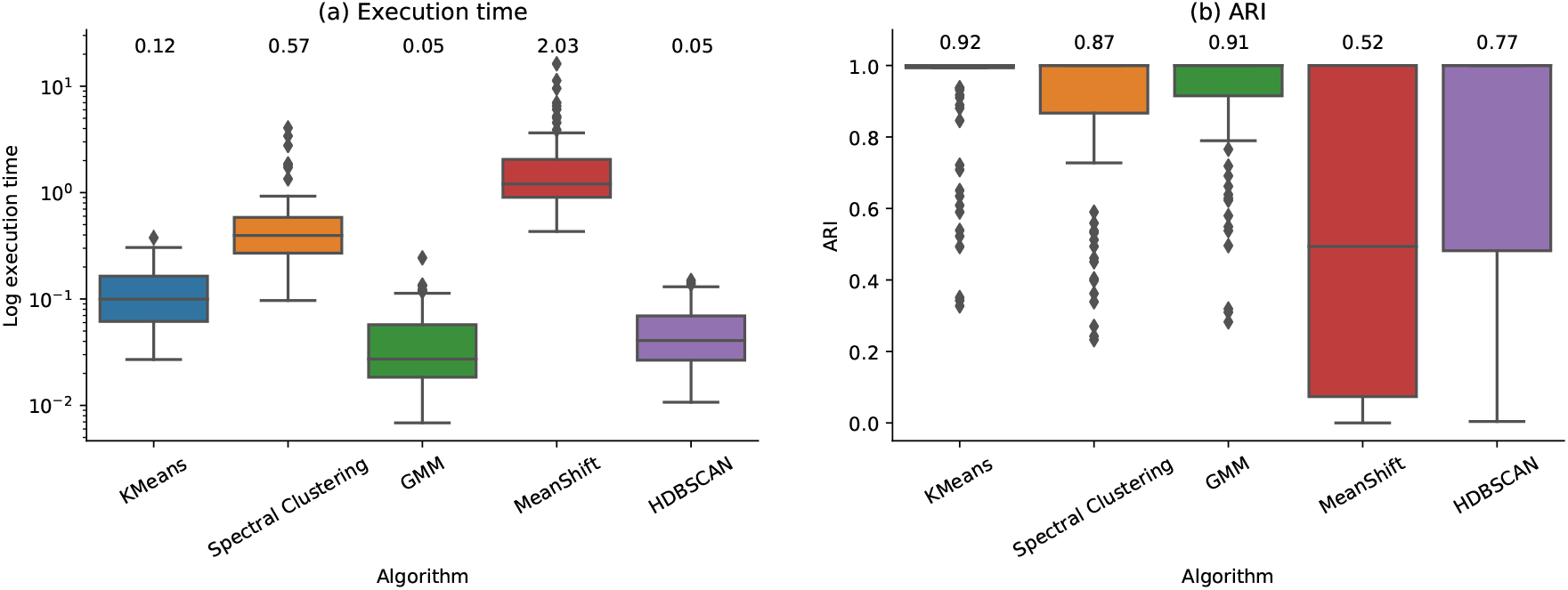
High level comparison between selected clustering algorithms with respect to the execution time expressed in seconds (a) and the Adjusted Rand Index (b). The values depicted at the top of each boxplot represent the average score per clustering algorithm. For the algorithms dependent on the input number of clusters (i.e. Kmeans, GMM, Spectral Clustering), GMM is the fastest and has performances very close to Kmeans. For the clustering algorithms dependent on data density (i.e. MeansShift and HDBSCAN), HDBSCAN brings the fastest and the closest to the ground truth results. The performances of density-based algorithms are generally lower than that of algorithms working with a predefined number of clusters.

### A.11 Importance of individual sampling techniques

This section illustrates the relative importance of each feature sampling technique and their impact on the subspace clustering performances measured with ARI score and percentage of identified features. As a reminder, the feature sampling consists of identifying important features, abbreviated as I (using the uni dimensional feature ranking), the proximal features to the subspace to optimize, abbreviated as P and random feature exploration, abbreviated as R.

Each dataset is analyzed using the complete set of combinations between the three feature sampling strategies. Thus, after running our method using only one of the three sampling methods (I, P, R), configurations consisting of combining two and all sampling methods with equal probability are tested. The results in Figure 19a depict the average ARI score on the two identified subspaces and indicate that combining important and proximal features produces the most accurate results (average ARI score of 93), followed by the combination including random exploration (average ARI score pf 0.90). The results in Figure 19b depict the percentage of identified features and suggest that, the random exploration has generally a positive effect on the completeness of identified subspaces. The highest score corresponds to the combination of all sampling strategies (I, P, R) with equal probability. This setting reflects also the default configuration of our method used when generating all experimental results presented previously. As expected, the worst performance is reported when employing only the random exploration, results which confirm empirically the importance of the proposed feature sampling method.

**Fig. 19.**
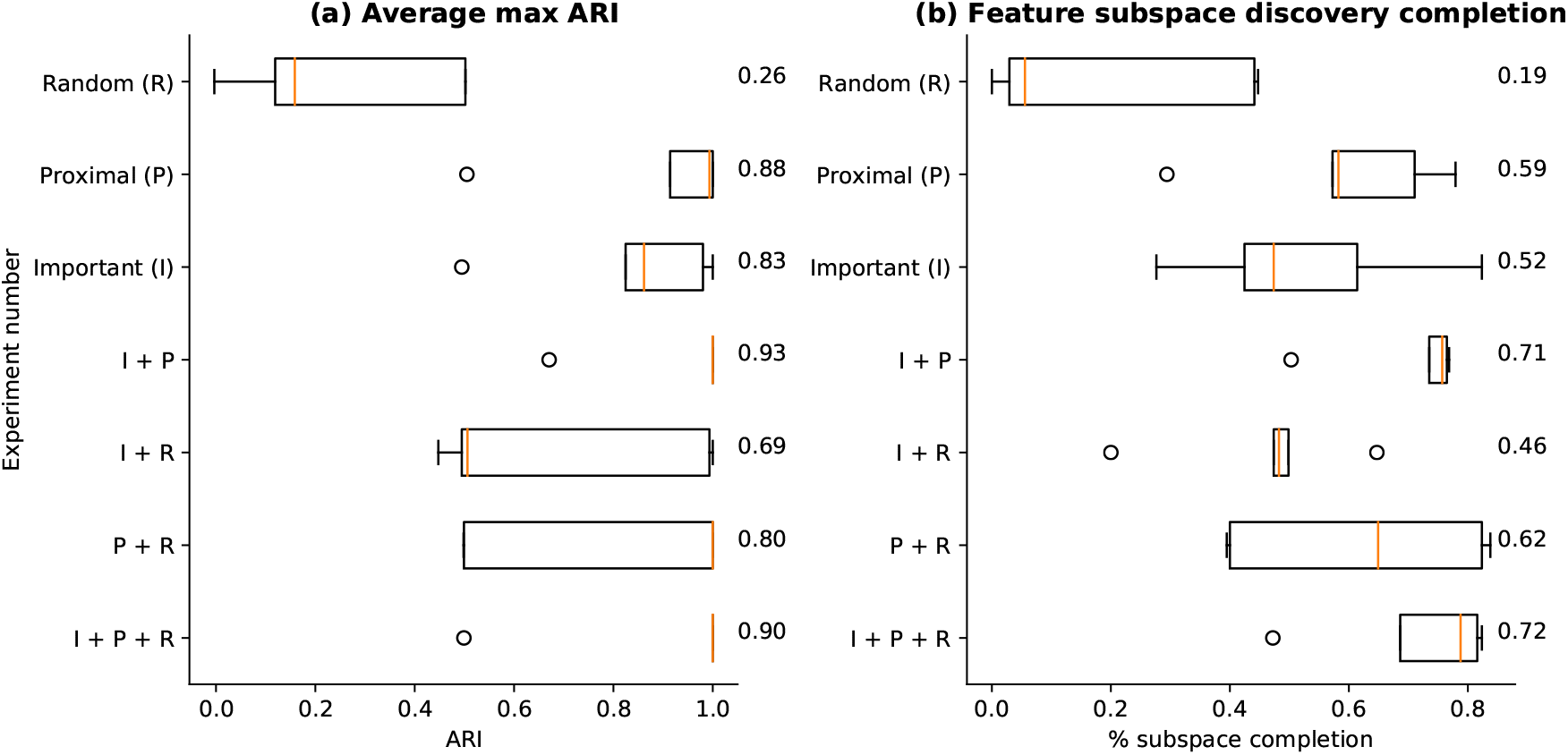
Importance of each feature sampling method for the average subspace ARI score (panel a) and the average subspace feature identification rate (panel b). A set of 5 simulated datasets are generated to contain 2 subspaces and noisy features. Each dataset is analyzed with our method using all 7 combinations of feature sampling methods: Important features (I), proximal features to the subspace to optimize (P), random exploration (R) and all permutations of 2 and 3 methods (i.e. I + P, I + R, P + R, I + P + R). The experimental results suggest that addition of important and proximal features to the random exploration provides a significant performance improvement.

### A.12 Strategies to interpret the discovered subspaces on RNA-seq datasets

This section presents several strategies to interpret the discovered subspaces, by using the patient metadata, annotated cancer genes or by performing survival and cell enrichment analysis.

#### A.12.1 Patient metadata

Patient metadata is only available for the RNA-seq datasets and provides information concerning the vital status, the number of days to death, the gender, the age and annotations corresponding to the presence of the disease and other pathology-specific indicators. The relation between the discovered subspaces and all other annotated criteria has been evaluated by computing ARI scores. First, we analyzed the results wrt the disease subtype annotation. On BRCA dataset ( Figure 10a), a subspace has been identified having a 0.68 ARI score wrt the ground truth, which is close to the score leveraging top 10 features selected in a supervised way (0.71). On the KIRP dataset ( Figure 10b) a subspace having an ARI score of 0.17, while top 10 features selected in a supervised way have a maximim ARI score of 0.38.

Next, the patient metadata is explored in a similar way, considering each annotated trait as a new ground truth variable and computing the underlying ARI score. For each one of our subspaces, we computed the ARI scores with respect to each annotation. The results in Figure 10 depict for each subspace, the annotated trait having the highest ARI score. Even though the disease indicator has no missing values, on average half of this information is not present. For this reason, the ARI score has been computed only on the observations present in the annotation. Figure 10 shows that on the KIRP dataset a subspace corresponding to gender has been identified (0.97) followed by the clinical Stage (0.4). BRCA has less well defined clusters; the disease related subspace has been identified (0.68), as well as the BCR Canonical reason (0.46).

#### A.12.2 Survival analysis

Another perspective to assess the results is by searching for relations between the patient partitions, identified in each output subspaces, and their survival rate. For each subspace, the Kaplan-Meier curves estimating the fraction of patients living for a certain amount of time after treatment have been computed. Additionally, for each subspace the log-rank test has been performed, a non-parametric statistic used to compare the survival distributions of two samples. As depicted in Figure 20 two significant subspaces are identified for the BRCA dataset, best matching the Canonycal Reason and the Estrogen Receptor, while for KIRP the significant subspace corresponds to the Lactate dehydrogenase. However, not finding a significant correlation between the identified partitions and the survival rate does not invalidate the quality or the importance of discovered subspaces. The survival function is a complex topic, subject to both genetic and external factors; furthermore patient distinctions may not always manifest in aligned survival rates.

**Fig. 20.**
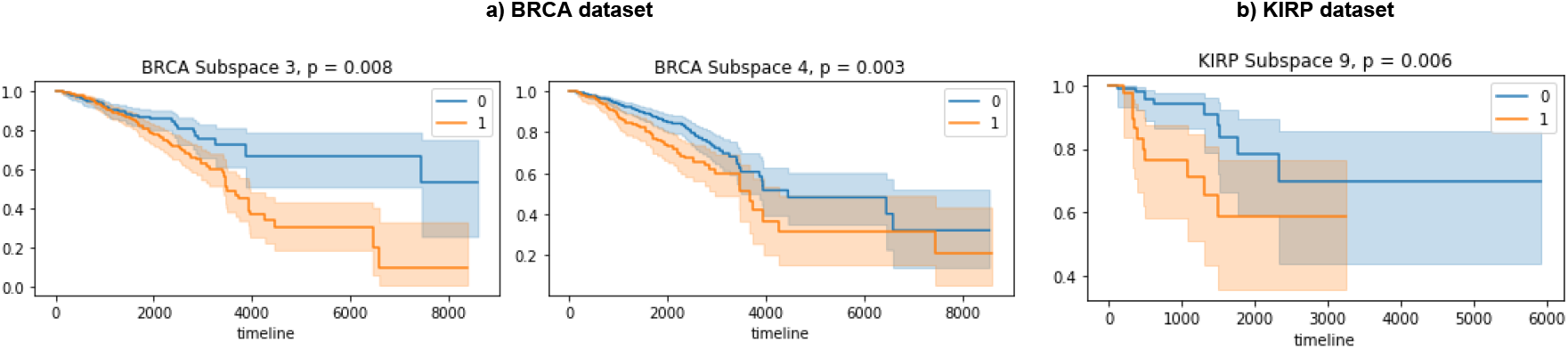
Illustration of survival curves on BRCA (panel a) and KIRP (panel b) datasets for the subspaces with a logrank test score < 0.05

**Fig. 21.**
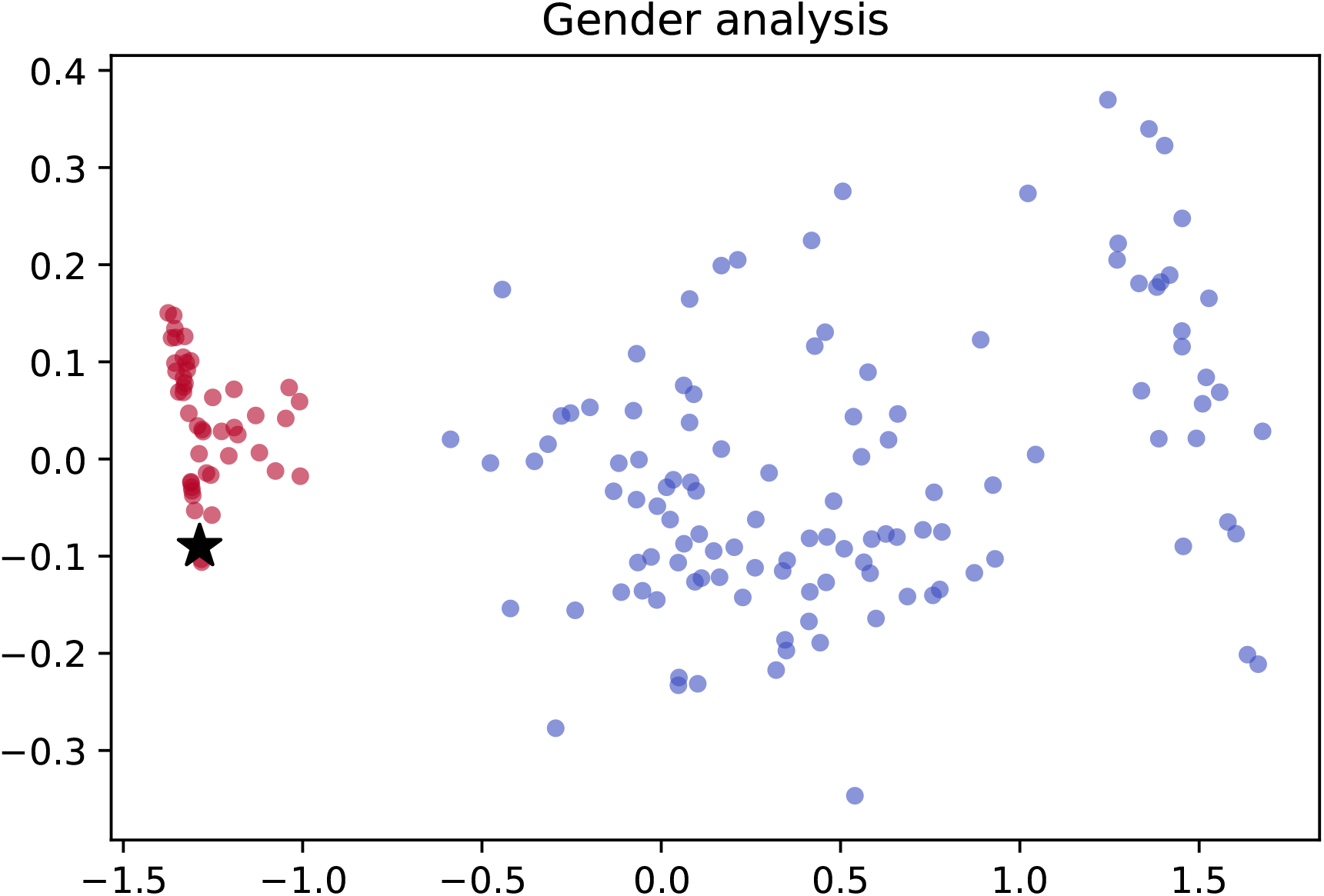
Example of one subspace discovered on the KIRP dataset which corresponds to gender separation (ARI = 0.97). The 2D representation has been obtained with PCA and plotting the first two components. The colored clusters (red and blue) correspond to the predicted classes. Only one patient (depicted by the black star) has been mislabeled by the algorithm, as the ground truth annotation places it in the blue cluster. We hypothesize that this is likely an annotation error.

#### A.12.3 Cancer genes analysis

Since both RNA-seq datasets analyze cancer patients, we studied the overlap between the discovered subspace features and documented cancer genes. If a discovered subspace contains a high percentage of genes known as markers for a certain condition, we have reason to investigate further its link in the context of this disease. A compilation of multiple studies such as Atlas ^9^, Sanger ^10^ and others ^11^ has been employed as reference. Our analysis shows that more than half of the discovered subspaces contain cancer genes, as detailed in Table 4. This exercise provide an alternative strategy for the biological feature-wise validation of results.

**Table 4.**
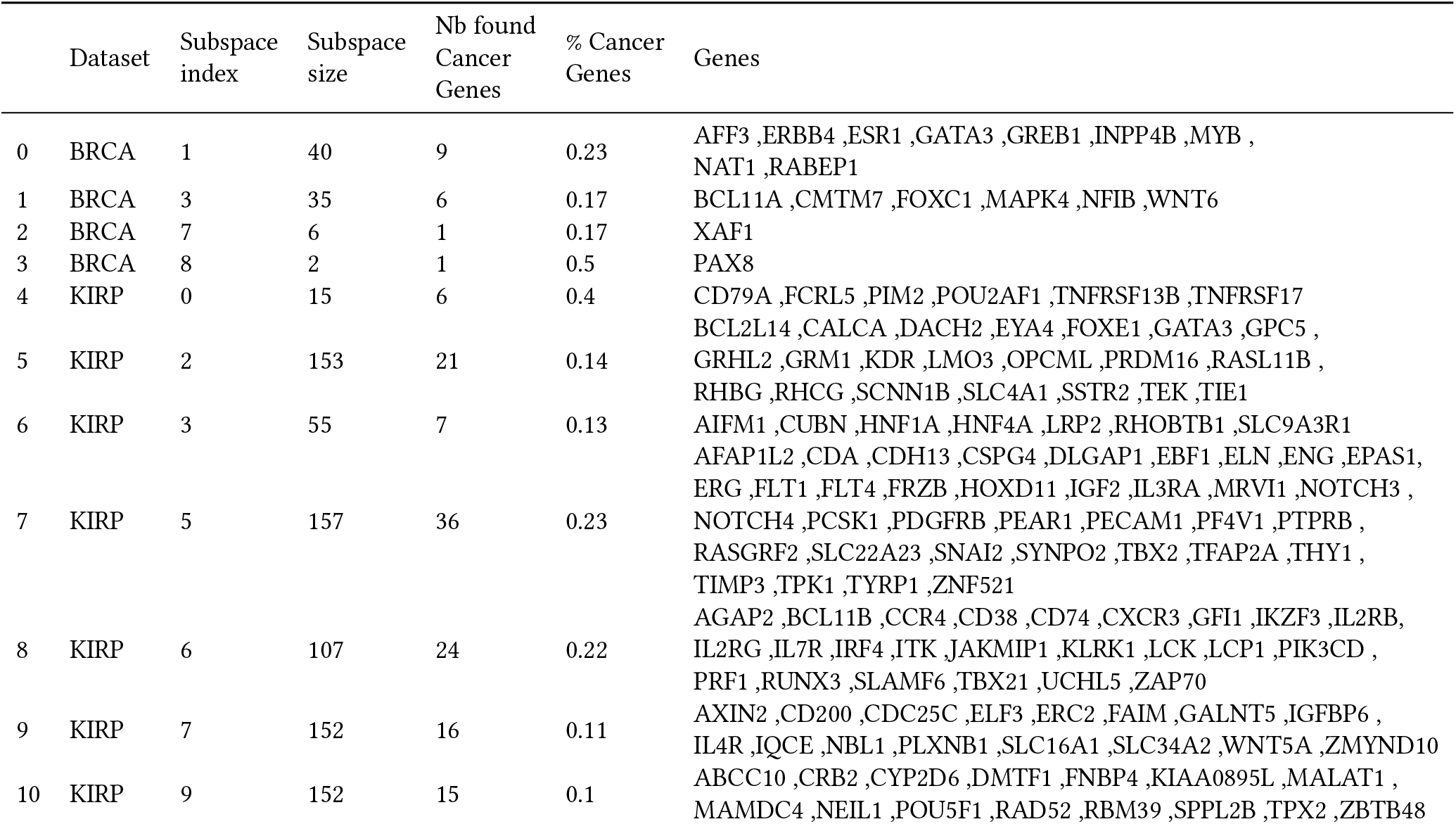
Documented Cancer Genes present in each of the discovered subspaces on BRCA and KIRP datasets. Each dataset has more than one subspace composed by a significant number of cancer genes. This is an alternative strategy for a feature-wise validation of results.

#### A.12.4 Gene ontology enrichment analysis

We propose an alternative method to interpret the significance of the discovered subspaces using gene ontology data. Gene ontology (GO) is a formal representation of the knowledge on molecular functions, cellular components and biological processes ^12^. We leverage this information to determine for each subspace if the underlying genes are coherent form a biological point of view and have a common functional link. Thus, we assess if there is an ontology of known cellular processes for which the genes in a given subspace represent an enrichment.

We employed the 2018 version of GO assembled through a python package ^13^, and we tested the discovered subspace on BRCA and KIRP datasets using the biological processes, cellular components and molecular functions libraries. In order to avoid spurious correlations, we selected for testing the subspaces having more than 10 features. The results in the table below show that for both datasets have been identified Molecular Functions with an adjusted p-value<0.05. For BRCA 3 feature subspaces are retrieved. The most important function names by p-value are depicted in Table 5. Four significant subspaces have been selected from the KIRP dataset, having on average 6 function values.

**Table 5.** Gene Ontology results on BRCA and KIRP datasets. We tested all subspaces for biological processes, cellular components and molecular functions. For each dataset we found at least one significant subspace. In both cases, multiple molecular functions have been identified.

| Dataset | Tested GO Libraries | Nb tested vs discovered subspaces | Nb significant subspaces and their functions |
| --- | --- | --- | --- |
| BRCA | GO_Biological Process 2018,<br>GO_CellularComponent2018,<br>GO_MolecularFunction 2018 | 3 / 10 | 1 with 13 Molecular Functions |
| KIRP | GO_Biological Process 2018,<br>GO_CellularComponent2018,<br>GO_MolecularFunction 2018 | 5 / 10 | 4 having only Molecular Functions,<br>on average 6 different values per subspace |

However, without more in depth information about relevant functions or processes, interpreting the significance of the discovered subspaces remains a difficult task to due to the current incomplete understanding of all genes and their functions. Nevertheless, this interpretation strategy may prove valuable in settings leveraging prior knowledge about the researched processes.

### A.13 Stability across consecutive runs

The addition of random exploration to our experiments allows the method to produce two different solutions as a result of consecutive run. In this section, the stability across consecutive runs is measured by executing our method twice on biological datasets. For simplicity, four microarry and bulk RNA-seq datasets have been selected for a detailed analysis. The overlap between the subspace features of the two runs is computed ( Figure 22a), as well as its statistical significance using the hyper-geometric overlap score. The statistical scores reported in Figure 22b suggest a significant overlap between the subspaces identified across the consecutive runs.

**Fig. 22.**
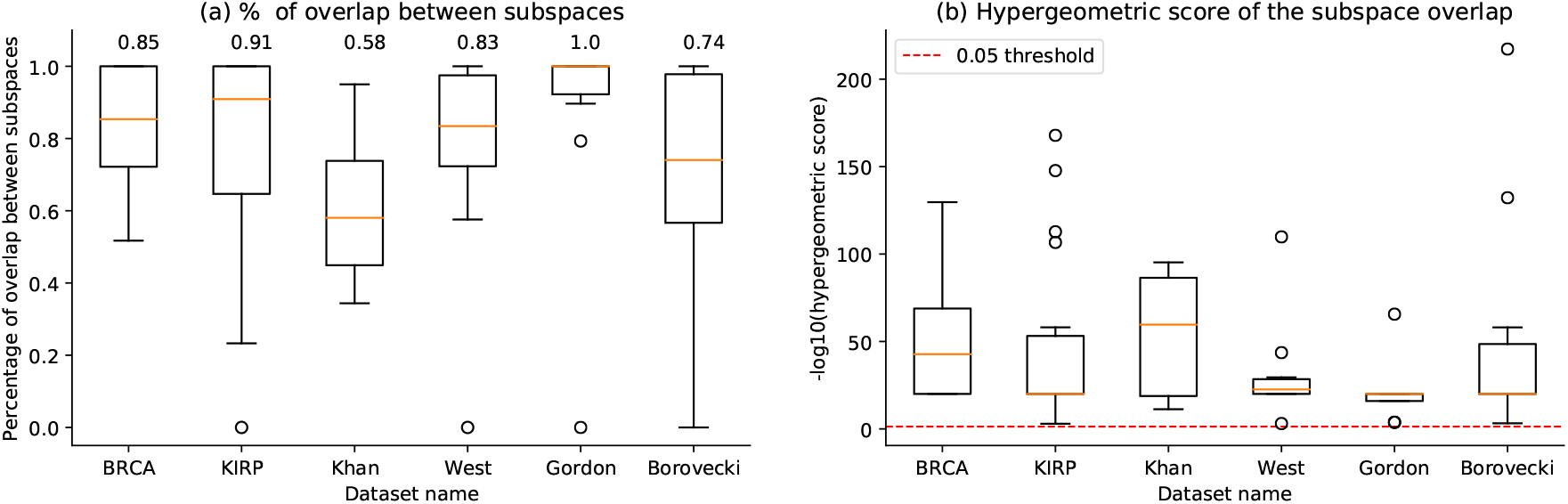
Assessment of the result determinism of the results in terms of the subspace features selected over consecutive runs of the method. We computed the number of features (expressed in percentage) that overlap in top 10 subspaces (a) and the statistical score of the overlap (b) computed using the hypergeometric distribution. On average, more than 50% of each subspace is reconstructed with a new run, depending on the statistical properties of the input datasets but also on the arbitrary percentage of random exploration.

### A.14 Semi-supervised analysis

Figure 23 summarizes the results of the comparative analysis between the unsupervised and the semi-supervised modes of our method.

**Fig. 23.**
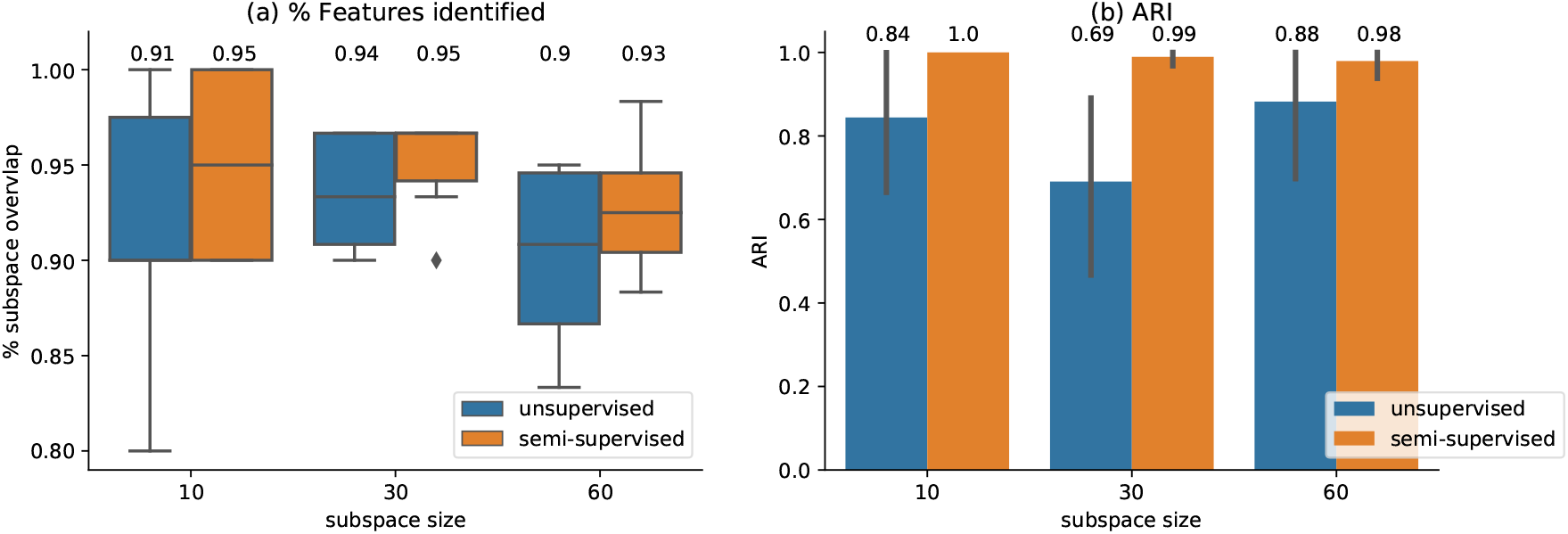
Semi-supervised versus unsupervised exploration. (a) Percentage of subspace overlap between ground truth embedded subspaces of various sizes and the results of the optimization algorithm running in unsupervised (blue) and semi-supervised (orange) modes. (b) ARI scores of all analyzed subspaces in unsupervised (blue) and semi-supervised (orange) modes. This analysis has been performed on simulated data. In all explored scenarios, the semi-supervised mode outperforms the unsupervised setting.

### Footnotes

1 https://scikit-learn.org/stable/

2 https://naeglelab.github.io/OpenEnsembles/

3 https://hdbscan.readthedocs.io/en/latest/

4 https://www.cancer.gov/about-nci/organization/ccg/research/structural-genomics/tcga

5 https://github.com/ramhiser/datamicroarray

6 https://rdrr.io/github/ramhiser/datamicroarray/

7 https://www.cancer.gov/about-nci/organization/ccg/research/structural-genomics/tcga

8 https://naeglelab.github.io/OpenEnsembles/

9 http://atlasgeneticsoncology.org

10 http://www.sanger.ac.uk/genetics/CGP/Census/

11 http://www.bushmanlab.org/links/genelists

12 http://geneontology.org/

13 https://pypi.org/project/gseapy/

## Notes

### Competing Interest Statement

The authors have declared no competing interest.

